# A model for Scc2p Stimulation of Cohesin’s ATPase and its Inhibition by Acetylation of Smc3p

**DOI:** 10.1101/2022.11.23.517716

**Authors:** Kevin M. Boardman, Siheng Xiang, Fiona Chatterjee, Udochi Mbonu, Vincent Guacci, Douglas Koshland

**Affiliations:** Department of Molecular and Cell Biology, University of California, Berkeley, Berkeley, CA 94720, USA

## Abstract

The evolutionarily conserved cohesin complex mediates sister chromatid cohesion and facilitates mitotic chromosome condensation, DNA repair, and transcription regulation. These biological functions require cohesin’s two ATPases, formed by the Smc1p and Smc3p subunits. Cohesin’s ATPase activity is stimulated by the Scc2p auxiliary factor. This stimulation is inhibited by Eco1p acetylation of Smc3p at an interface with Scc2p. It was unclear how cohesin’s ATPase activity is stimulated by Scc2p, or how acetylation inhibits Scc2p, given that the acetylation site is distal to cohesin’s ATPase active sites. Here, we identify mutations in budding yeast that suppressed the in vivo defects caused by Smc3p acetyl-mimic and acetyl-defective mutations. We provide compelling evidence that Scc2p activation of cohesin ATPase depends upon an interface between Scc2p and a region of Smc1p proximal to cohesin’s Smc3p ATPase active site. Furthermore, substitutions at this interface increase or decrease ATPase activity to overcome ATPase modulation by acetyl-mimic and - null mutations. Using these observations and a cryo-EM structure, we propose a model for regulating cohesin ATPase activity. We suggest that Scc2p binding to Smc1p causes a shift in adjacent Smc1p residues and ATP, stimulating the Smc3p ATPase. This stimulatory shift is inhibited through acetylation of the distal Scc2p-Smc3 interface.

## Introduction

The evolutionarily conserved protein complex called cohesin mediates sister chromatid cohesion and facilitates mitotic chromosome condensation, DNA repair, and transcription regulation. Cohesin is thought to perform these remarkably diverse biological functions through the complex control of its two activities, tethering two chromatin regions together (within or between DNA molecules) or extruding chromatin loops. Elucidating the different mechanisms of cohesin regulation and their coordination remains an important but elusive goal. Here, our studies in Saccharomyces cerevisiae provide a molecular mechanism for the coordinated control of cohesin by two of its key regulators, Scc2p and Eco1p.

Cohesin’s core complex contains four subunits, which in budding yeast are called Smc1p, Smc3p, Scc3p, and Mcd1p (Scc1p). Cohesin has ATPase activity. This activity requires two active sites (Smc3 ATPase and Smc1 ATPase) that are formed through the heterodimerization of the Smc1p and Smc3p head domains (Supplemental Fig. 1A) (Weitzer et al. 2003; Arumugam et al. 2003). Both active sites are required for cohesin’s ATPase activity, its loading onto chromosomes and all of its biological activities (Weitzer et al. 2003; Arumugam et al. 2003). In addition to the core complex, a heterodimer of Scc2p and Scc4p is required for cohesin to bind chromosomes and extrude loops (Ciosk et al. 2000; Bauer et al. 2021; Davidson et al. 2019). These activities are thought to derive from Scc2p’s ability to stimulate cohesin’s ATPase activity (Murayama and Uhlmann 2014; Petela et al. 2018; Çamdere et al. 2015). Recent Cryo-EM structures of yeast and human cohesin with Scc2p and its human ortholog, NIPBL, show that Scc2p has multiple interactions with cohesin’s head domains and Smc3p’s coiled-coil (Shi et al. 2020; Collier et al. 2020). The presence of these interfaces strongly suggests that Scc2p directly regulates the cohesin head domain and its ATPase activity.

**Figure 1:**
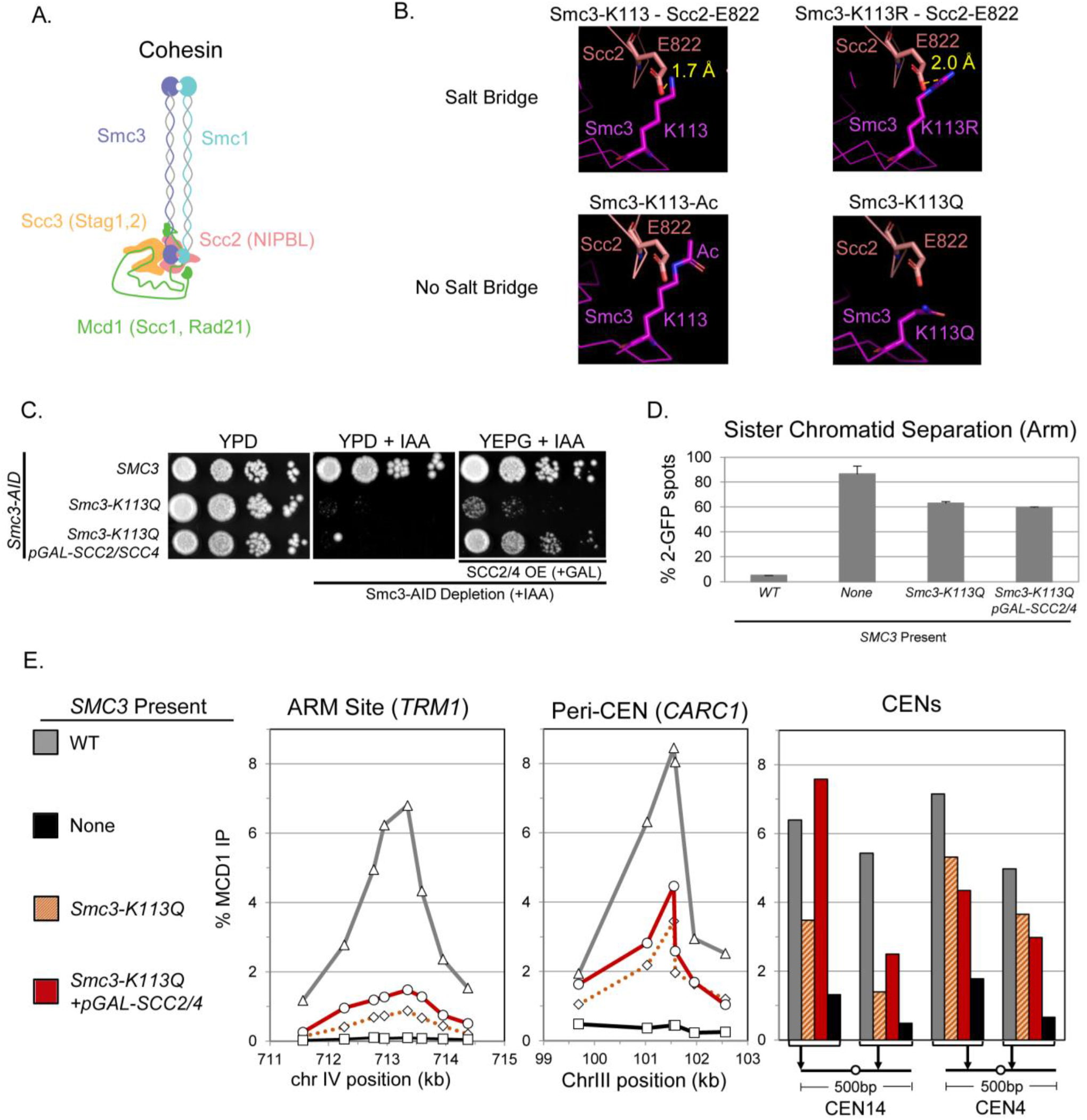
Characterization of the Acetyl-mimic and its Suppression by Scc2p/Scc4p Overexpression. **(A)** Cartoon depiction of cohesin (Smc1p, Smc3p, Scc3p, Mcd1p) with the associated stimulatory factor Scc2p (NIPBL). **(B)** Cryo-EM of S. cerevisiae cohesin illustrating that Smc3p-K113 acetylation eliminates a salt bridge with Scc2p-E822. Top panels show that salt-bridge can form between Smc3p-K113 (left) or K113R mutation (right) and Scc2p-E822 due to close proximity. Bottom panels illustrate that Smc3p-K113p acetylation (left) or K113Q mutation (right) alters spacing, precluding a salt bridge. Smc3p-K113 (magenta), Scc2p-E822 (salmon) and distance for putative salt-bridge (yellow). **(C)** SCC2/SCC4 overexpression suppresses the inviability of the smc3-K113Q acetyl-mimic. Haploid SMC3-AID strain also bearing either a wild-type SMC3 (VG3919-3C), smc3-K113Q (KB62A), or smc3-K113Q containing pGAL-SCC2/SCC4 (VG4052-3A), were grown at 30°C to saturation, plated at 10-fold serial dilutions on YPD, YPD + IAA, or YEPG+ IAA and incubated for 4 days at 23°C. **(D)** SCC2/SCC4 overexpression fails to suppress the cohesion defect of smc3-K113Q cells. Strains in (C) along with a haploid containing SMC3-AID as the sole SMC3 (VG3651-3D) were arrested in G1, depleted for SMC3-AID, SCC2/SCC4 overexpressed then synchronously released from G1 and arrested in mid-M phase. Cells were fixed and processed to score cohesion at a chromosome IV arm locus using the LacO-LacI system as described in Materials and Methods. The number of GFP spots was scored in mid-M phase arrested cells and the percentage of cells with defective cohesion (2 GFP spots) plotted. 200 cells were scored for each data point and data generated from two independent experiments. **(E)** Cohesin binding to chromosomes is greatly reduced in smc3-K113Q cells and only slightly increased by SCC2/SCC4 over-expression. Aliquots of mid-M phase cells from (D) were fixed and processed for ChIP using anti-Mcd1p antibodies as described in Materials and Methods. Mcd1p binding was assessed by qPCR and presented as a percentage of input DNA. Left panel is chromosome IV arm CAR region (TRM1), middle panel is chromosome III peri-centric region (CARC1), right panel are regions immediately flanking CEN14 and CEN4.

Cohesin is also regulated through the acetylation of its subunits by the Eco1p acetyltransferase. Eco1p acetylates two conserved lysines in the Smc3p head domain, which occurs after cohesin binds DNA in S phase (Supplemental Fig. 1) (Unal et al. 2008; Rolef Ben-Shahar et al. 2008; Zhang et al. 2008). In Saccharomyces cerevisiae, these lysines are at positions 112 and 113. Substitution of arginine for K113 (K113R) or both K112 and K113 (K112R, K113R) prevents their acetylation, causing inviability and a failure to establish sister chromatid cohesion (Unal et al. 2008; Rolef Ben-Shahar et al. 2008). Eco1 mutants have the same defects as the smc3-K112R, K113R, indicating these two Smc3p lysines are key targets of Eco1p. Likewise, the viability of smc3-K112R, K113R and eco1Δ mutants can each be restored by deleting the WPL1 gene (Rolef Ben-Shahar et al. 2008; Sutani et al. 2009; Rowland et al. 2009). Wpl1p is a conserved cohesin inhibitor that removes cohesin from DNA (Kueng et al. 2006; Chan et al. 2012). Thus, one evolutionarily conserved function of K113 acetylation is to antagonize Wpl1p and stabilize cohesin’s binding to DNA.

However, WPL1 deletion minimally suppresses the sister chromatid cohesion defects of either smc3-K112R, K113R or eco1Δ mutants (Rowland et al. 2009; Sutani et al. 2009; Guacci and Koshland 2012; Guacci et al. 2015). Other mutations have been identified that do suppress the cohesion defects of acetyl-defective mutants (Çamdere et al. 2015; Elbatsh et al. 2016)(Rowland et al. 2009; Sutani et al. 2009; Guacci and Koshland 2012; Guacci et al. 2015)(Çamdere et al. 2015; Elbatsh et al. 2016). These mutations map to the Smc3 ATPase active site and reduce cohesin ATPase activity (Çamdere et al. 2015; Elbatsh et al. 2016). This observation suggests that cohesin’s unacetylated and acetylated states impose different cohesin ATPase levels. Initially, the higher ATPase levels of unacetylated cohesin induced by Scc2p promote efficient cohesin DNA binding. Once cohesin is bound to a sister chromatid, Smc3p acetylation lowers cohesin’s ATPase activity. The lower ATPase activity stabilizes its DNA binding and promotes the capture of the other sister chromatid to generate cohesion (Çamdere et al. 2015; Elbatsh et al. 2016); (Çamdere et al. 2018). The mutants lacking Smc3p acetylation cannot stably bind DNA or capture a second DNA molecule because its ATPase activity is too high. This defect of smc3-K112R, K113R or eco1Δ mutants is counteracted by the suppressor mutations in the Smc3 ATPase active site that slow ATP hydrolysis (Çamdere et al. 2015, 2018).

How acetylation alters cohesin’s ATPase has remained a mystery as the Smc3p-K113 residue is not proximal to either of cohesin’s ATPase active sites (Supplemental Fig. 1A). A recent study suggested that Smc3p-K113 acetylation inhibited Scc2p binding to cohesin indirectly by stabilizing the binding of Pds5p, which then precluded Scc2p binding (Bastié et al. 2022; van Ruiten et al. 2022). However, a more direct mechanism was suggested by the observation that Scc2p stimulation of cohesin ATPase activity in vitro is drastically reduced by a mutation that mimics acetylation by substituting glutamine for K113 (K113Q) (Murayama and Uhlmann 2015). Notably, this inhibition occurred in the absence of Pds5p, suggesting acetylation controls cohesin ATPase by a second Pds5p-independent mechanism.

Here, we test and elaborate on this hypothesis by addressing three mechanistic questions: 1) Does acetylating Smc3p-K113 impede Scc2p function in vivo, as suggested by the in vitro experiments? 2) How does Scc2p activate cohesin’s ATPase? 3) How is cohesin’s ATPase activity repressed by acetylation of the Smc3-K113 residue, given that this residue lies far away from either of cohesin’s ATPase active sites? The answers to these questions led us to a model that explains both Scc2p’s activation of cohesin’s ATPase and its inhibition by acetylation.

## Results

### Inhibiting the salt bridge between Smc3-K113 and Scc2p compromises cohesin function in vivo

Further clues into how Smc3p-K113 acetylation could impact Scc2p function came from the cryo-EM structures of yeast and human cohesin with Scc2p or its human ortholog, NIPBL (Collier et al. 2020; Shi et al. 2020). The Smc3p-K113 residue and its homologous residue in humans, Smc3p-K106 (Zhang et al. 2008), are positioned to make a salt bridge with the carboxyl group of conserved Scc2p glutamate (E821 in Scc2p and E1899 in human NIPBL) (Figure 1B, Supplemental Fig. 1B-C). This salt bridge suggested that the K113 residue directly contributed to Scc2p’s stimulation of cohesin’s ATPase by facilitating Scc2p binding to the cohesin head.

The involvement of the Smc3p-K113 residue in this salt bridge provided a simple mechanism for how its acetylation, the acetyl mimic (smc3-K113Q), and the acetyl-defective (smc3-K113R) could alter Scc2p’s stimulation of the cohesin ATPase. The positively charged guanidino nitrogens of the K113R substitution should still be able to form the salt bridge with the negatively charged carbonyl oxygen of the Scc2p-E821 residue. To test this assumption, we made the arginine substitution in the cryo-EM structure. Indeed, the guanidino nitrogens were close enough (<4 angstroms) to form an effective salt bridge and should allow the binding of Scc2p and its stimulation of the cohesin ATPase (Figure 1B). However, since the arginine substitution cannot be acetylated, the salt bridge should persist constitutively, causing the smc3-K113R mutant to phenocopy eco1 mutants (Unal et al. 2008; Rolef Ben-Shahar et al. 2008; Guacci and Koshland 2012).

Conversely, acetylation of the Smc3p-K113 residue or the smc3p-K113Q substitution would eliminate the lysine’s positive charge and block the formation of the salt bridge with Scc2p. The disruption of the salt bridge would alter Scc2p’s binding to cohesin such that it could no longer stimulate the ATPase (Shi et al. 2020; Murayama and Uhlmann 2015). While K113 acetylation and smc3p-K113Q would block the salt bridge, they would not generate structural conflicts that would dramatically impact the cohesin head structure (Figure 1B).

To better understand the impact of Smc3p-K113 acetylation, we performed a detailed characterization of the smc3-K113Q mutant. For this purpose, we tagged the endogenous SMC3 gene with an auxin-inducible degron (SMC3-AID). Then, we introduced a second SMC3 allele, wild-type SMC3 or smc3-K113Q. The addition of auxin (IAA) to the growth media for these cells induced the degradation of the Smc3p-AID protein, generating cells either lacking Smc3p (SMC3-AID) or containing only wild-type Smc3p (SMC3-AID SMC3), or only smc3p-K113Q (SMC3-AID smc3-K113Q). Cells lacking Smc3p or containing only smc3p-K113Q were inviable (Figure 1C), corroborating a previous study that smc3-K113Q disrupts an essential cohesin function (Heidinger-Pauli et al. 2010).

To further characterize the smc3-K113Q mutant, we analyzed cells from these three strains for sister chromatid cohesion at a centromere-proximal and arm locus by fluorescently marking each sister chromatid with the LacO-lacI-GFP system (Marshall et al. 1997; Guacci et al. 2019). We depleted Smc3p-AID from G1 through M phase and assessed sister chromatid cohesion in mid-M phase-arrested cells. The precocious sister chromatid separation was detected by the presence of two GFP spots. The percentage of cells with two spots increased dramatically in cells expressing only smc3p-K113Q compared to Smc3p, increasing 3 to 4-fold near the centromere and10-fold at the chromosome arm locus (Figure 1D, Supplemental Fig. 1D). These cohesion defects were not as severe as in cells containing no Smc3p (Figure 1D, Supplemental Fig. 1D). Thus, the acetyl mimic mutant was severely but not completely defective for sister chromatid cohesion, as seen in previous studies (Unal et al. 2008; Guacci and Koshland 2012).

Cohesin is bound to chromosomes at the centromeres, pericentric regions, and specific sites on chromosome arms from S phase to mid-M. To assess how Smc3p-K113Q affects cohesin binding to DNA, we performed chromatin immunoprecipitation followed by quantitative PCR (ChIP-qPCR) on aliquots of synchronously arrested cells in mid-M phase. Cells containing only Smc3p-K113Q showed a 6-fold reduction in cohesin binding to an arm site, a 2-fold decrease at a pericentromeric site, and a variable loss of binding at centromeres compared to cells containing Smc3p (Figure 1E). However, the Smc3p-K113Q cohesin binding levels at all these sites were well above those observed in cells lacking Smc3p, indicating that DNA binding of cohesin was significantly but not entirely compromised by Smc3p-K113Q. Taken together, the multiple defects of cells containing only Smc3p-K113Q were consistent with the idea that blocking the salt bridge between Smc3p-K113 and Scc2p inhibits Scc2p function and presumably its ability to stimulate cohesin ATPase.

Finally, we examined whether Smc3p-K113Q compromised the integrity of the cohesin core complex by assessing the levels of the Mcd1p subunit of cohesin in mid-M arrested cells. Mcd1p is degraded unless associated with both Smc1p and Smc3p (Çamdere et al. 2015; Guacci et al. 2015; Robison et al. 2018; Guacci et al. 2019). Mcd1p levels were reduced 4-fold in Smc3p-K113Q cells compared to Smc3p but remained at least 2-fold higher than those lacking Smc3p (Supplemental Fig. 1E). This result suggested that the acetyl-mimic mutant partially destabilized Mcd1p’s binding to cohesin.

Having completed this detailed characterization of the smc3-K113Q mutant, we assessed whether any of its phenotypes were suppressed by Scc2p/Scc4p overexpression as predicted if the acetyl-mimic mutation partially compromised Scc2p function. The inviability of the strain expressing only Smc3p-K113Q was suppressed by the galactose-induced overexpression of Scc2p and Scc4p (Figure 1C). In contrast, Scc2p/Scc4p overexpression did not suppress the inviability of cells that were depleted for either Eco1p or the cohesion maintenance factor Pds5p (Supplemental Fig. 1F). These results are consistent with the hypothesis that Smc3p-K113 acetylation limited the function of Scc2p, but not other cohesin regulators.

However, Scc2p/Scc4p overexpression minimally suppressed the Smc3p-K113Q-induced defects in cohesion, chromosome binding, and Mcd1p levels (Figure 1D-E, Supplemental Fig. 1D-E). The smc3-K113Q mutant overexpressing Scc2p/Scc4p was also sensitive to benomyl (a broader measure of cohesin’s mitotic function) and camptothecin (a measure of cohesin’s function in DNA damage repair) (Supplemental Fig. 1G). The presence of these defects showed that Scc2p/Scc4p overexpression only partially restored cohesin function in the smc3-K113Q mutant. The inefficiency of suppression could be explained by the inability of Scc2p/Scc4p over-expression to restore normal DNA binding levels to cohesin with Smc3p-K113Q.

To better understand how the smc3-K113Q mutation impacts Scc2p regulation of cohesin, we conducted a screen in budding yeast for suppressor mutations that restored viability to cells expressing only Smc3p-113Q (Figure 2A). We hypothesized that these mutations would identify regions of cohesin that were responsive to Scc2p and/or Smc3p-K113 acetylation. We identified three suppressor mutations in SMC3 and three in SMC1 (figure 2B). All of the suppressor mutations were located in regions of the cohesin subunits that had conserved amino acid sequences (Supplemental Fig. 2A-B). We introduced each of the six suppressor mutations into the parental smc3-K113Q cells and also strains in an otherwise wild-type background (Materials and Methods). All six newly constructed double mutants were viable, demonstrating that our six mutations were responsible for the suppression of smc3-K113Q inviability. Thus, these mutations identified regions of the cohesin complex, which were functionally connected to an acetylated state of the Smc3p-K113 residue.

**Figure 2:**
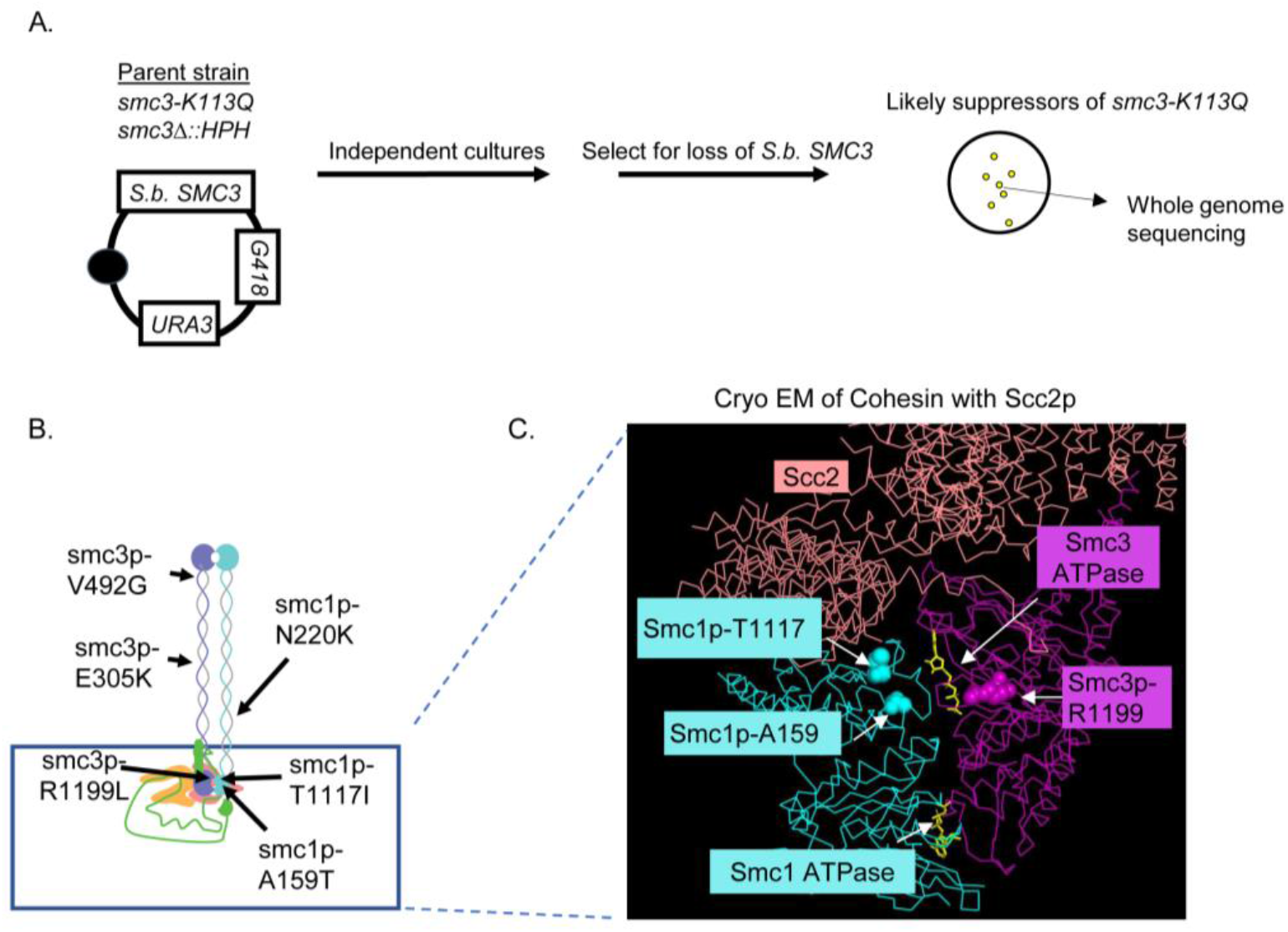
Suppressor screen links Scc2p-Smcp-K113 interface with other Scc2p-cohesin head interfaces. Identification of suppressor mutations in SMC1 and SMC3 which suppress the lethality of smc3-K113Q. **(A)** Schematic of the genetic screen used to identify smc3-K113Q suppressors. Haploid strain (VG3969-14C) bears smc3-K113Q as the sole S. cerevisiae copy of SMC3 and S. Bayanus SMC3 on a CEN URA3 G418 plasmid (pFC3). Single colonies were grown to saturation in YPD, plated on FOA media to select for viable cells expressing only smc3p-K113Q (see Materials and Methods). Representative colonies from each plate were sequenced to identify the putative suppressor mutation. **(B)** Cartoon of the cohesin complex showing relative positions of the smc3-K113Q suppressor mutations shown. Smc1p (cyan), Smc3p (purple), Mcd1p (green). Also depicted is cohesin-associating protein, Scc2p (salmon). **(C)** Residues of 2 suppressors, Smc1p-R1199 and Smc1-T1117 map close to the Smc3p ATPase and Scc2p. Cryo-EM structure of the S. cerevisiae Smc1p (cyan) and Smc3p (magenta) head domains bound to Scc2p (salmon). S. cerevisiae Smc3p-R1199 residue (magenta spheres), S. cerevisiae Smc1p-T1117 residue (cyan spheres), and S. cerevisiae Smc1p-A159 (cyan spheres) residues are indicated. ATP (yellow) and each SMC ATPase are also indicated.

We mapped our suppressor mutations onto the cryo-EM structure of cohesin with the Scc2p complex (Collier et al. 2020). The six suppressor residues lay in distinct regions of cohesin, three in the Smc coiled-coil domains and three in the Smc head domains (Figure 2B). This result suggested that multiple regions of cohesin were impacted by Smc3p-K113 acetylation (Figure 2B). The three suppressors in the head were much closer to the Smc3p ATPase active site than the Smc1p ATPase active site (Figure 2C). These results suggested that the acetylated state of Smc3p-K113 regulated cohesin by modulating the Smc3-ATPase active site.

Additional insights came from analyzing the local structure of the two residues that were mutated by the suppressors. The Smc1p-T1117 residue was proximal to an interface between Smc1p and Scc2p and this interface and positioning was conserved between yeast and humans (Supplemental Fig. 2C). The Smc3p-R1199 residue was missing from the yeast structure; however, this residue lay in a conserved region of Smc3p (Supplemental Fig. 2B). Using this conservation to map the orthologous Smc3p-R1199 region on the human structure revealed that it too was proximal to an interface between NIPBL (the Scc2p ortholog) and Smc3p (Supplemental Fig. 2D). While the Smc residues of both of these interfaces were conserved, the residues in NIPBL and Scc2p at these interfaces were not (Supplemental Fig. 2E-G), suggesting that both Scc2p interfaces were rapidly evolving. Taken together, the positions of the suppressor residues suggested that alterations in these Smc1p-Scc2p or Smc3p-Scc2p interfaces could act at a distance to compensate for the defects associated with the elimination of the Scc2p-Smc3p-K113 salt bridge by K113 acetylation. This compensation suggested that proper Scc2p function involved functional crosstalk between three distinct interfaces of Scc2p and the cohesin head.

To more fully evaluate the efficacy of these suppressors, we subjected strains with the suppressor mutation alone or with smc3-K113Q to the same battery of tests that we used to analyze suppression by Scc2p overexpression (Figure 1C-E, Supplemental Fig. 1E, 1G). Five of the six double mutants behaved similarly; they grew slowly, exhibited significant benomyl and camptothecin sensitivity, and only partially restored cohesion (Figure 3A-B, Supplemental Fig. 3A-D). The persistence of the phenotypes of these five mutants could have been caused by inefficient suppression of the smc3-K113Q allele, or new defects caused by the suppressor mutations. However, the suppressor mutations in an otherwise wild-type background had no obvious phenotypes (Supplemental Fig. 3A-D). Thus, the defects of the double mutants reflected the failure of these five suppressor mutations to fully suppress the defects in cohesin function caused by the smc3-K113Q mutation.

**Figure 3:**
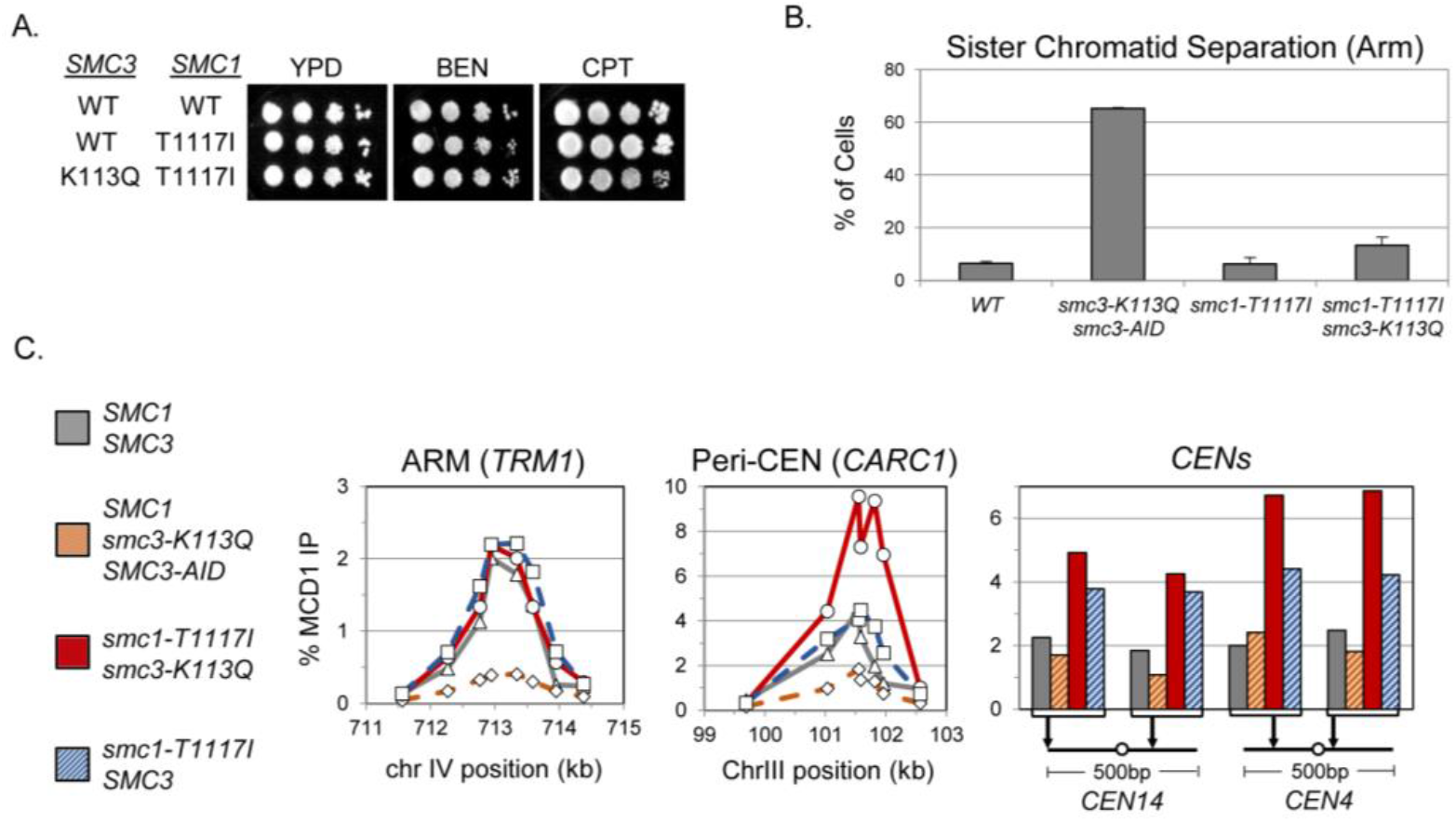
Smc1-T1117I uniquely and robustly suppresses smc3-K113Q defects in viability, drug sensitivity, cohesion and cohesin binding to DNA. The smc1-T1117I mutant is a robust suppressor of the smc3-K113Q mutant. **(A)** The smc1-T1117I smc3-K113Q double mutant grows as well as WT and is resistant to drugs. Haploid wild-type (VG4012-2C), smc1-T1117I (VG4006-13A) and smc3-K113Q smc1-T1117I (VG4010-8B), strains were grown and diluted as described in Figure 1C then plated on YPD alone or containing benomyl [BEN] (10mg/ml) or camptothecin [CPT] (15mg/ml) and incubated for 3d at 23°C, 4d at 23°C, or 3d at 30°C respectively. Plates were electronically rearranged for ease of display. **(B)** smc1-T1117I strongly suppresses the cohesion defect of the smc3-K113Q mutant. Haploid wildtype (VG3620-4C), smc3-K113Q smc3-AID double mutant (VG3891-6B), smc1-T1117I (VG4006-13A), and smc3-K113Q smc1-T1117I double mutant (VG4010-8B) cells were arrested in G1, auxin was added to deplete Smc3p-AID, then cells synchronously released from G1 and arrested in mid-M phase. Cohesion loss at a chromosome IV arm locus was assessed and plotted as described in Figure 1D. **(C)** The smc1-T1117I smc3-K113Q double mutant cohesin binds to chromosomes at levels equal to or higher than wild-type cohesin. Mid-M phase cells from (B) were fixed and processed for ChIP, then the level of cohesin bound to chromosomes determined as described in Figure 1E. Left panel: chromosome IV arm CAR region (TRM1); middle panel: chromosome III peri-centric region (CARC1); right panel: regions immediately flanking CEN14 and CEN4.

In stark contrast, the viability, growth, drug resistance, and cohesion of the smc1-T1117I smc3-K113Q double mutant strain were nearly indistinguishable from the wild-type (Figure 3A-B). Furthermore, the cohesin binding to DNA in this double mutant was completely restored to wild-type levels at an arm locus and elevated above wild-type levels at the pericentromeric and centromere regions (Figure 3C). Thus, this suppressor compensated for almost all the biological and molecular defects imposed by smc3-K113Q. The only defect not completely restored in the smc3-K113Q smc1-T1117I double mutant was that Mcd1p levels were still reduced compared to wild-tythe pe (Supplemental Fig. 3E-F). This result suggested that smc1-T1117I did not suppress smc3-K113Q mutant defects by simply increasing the level of mutant cohesin in cells. Rather the smc1-T1117I mutation must have improved the functionality of Smc3p-K113Q cohesin.

### A model for Scc2p activation of cohesin’s ATPase and its regulation by Smc3p-K113 acetylation

We reasoned that smc1-T1117I improved smc3-K113Q cohesin function by overcoming the acetylation-dependent inhibition of Scc2p stimulation of cohesin’s ATPase. A closer look at the cohesin-Scc2p cryo-EM structure provided an important clue as to how the suppressor mutation could impact ATPase activity. The Smc1p-T1117 residue lay immediately adjacent to arginine (R1122) and asparagine (N1096) residues in Smc1p (Figure 4A). These Smc1p residues likely contributed to Smc1p binding to Scc2p. Smc1p-R1122 was positioned to make a salt bridge with Scc2p’s glutamate at 1168 while Smc1p-N1096 was capable of hydrogen bonding with Scc2p’s threonine at 1210 (Figure 4A, Supplemental Fig. 4). The smc1p-T1117 residue was also immediately adjacent to the Smc1p-K1121 residue, which was positioned to form a hydrogen bond with a hydroxyl group on the ribose ring of ATP in the Smc3 ATPase active site (Figure 4A). Thus, the smc1p-T1117I suppressor identified a region of Smc1p that was positioned to both respond to Scc2p binding (through K1122 and N1096) and impact an Smc1p residue (K1121) important for ATP binding.

**Figure 3:**
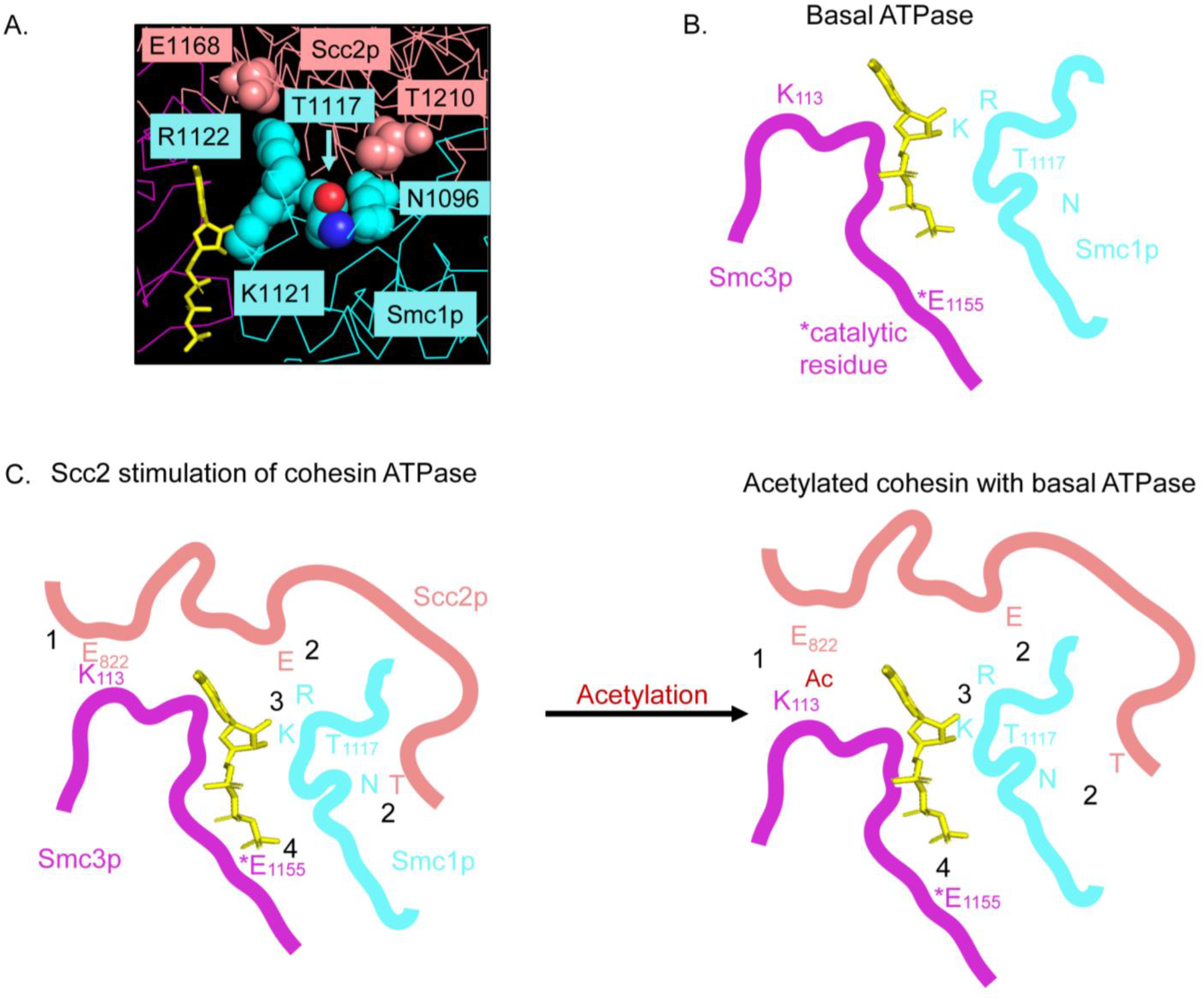
Model for Scc2p-mediated stimulation of Smc3p ATPase and its inhibition by acetylation. **(A)** Cryo-EM of the interface between Smc1p (cyan) and Scc2p (salmon), indicating Smc1-T1117 and other key Smc1p and Scc2p residues and ATP (yellow). **(B)** Cartoon of cohesin head domain in its basal ATPase state and with key Smc1p and Smc3p residues indicated. **(C)** Cartoon of cohesin head domain in its stimulated ATPase state with Scc2p bound when Smc3p in not acetylated (left), and the inhibited state with Smc3p-K113 acetylated (right). **(Left side)** Scc2p stimulation of cohesin ATPase: (1) Scc2p-E822 binds to unacetylated Smc3p-K113, which (2) properly orients Scc2p-E1168 (E) to interact with Smc1p-R1122 (R) and Scc2p-T1210 (T) to interact with Smc1p-N1096 (N). Scc2p binding at this Smc1p interface (3) shifts Smc1p and ATP such that (4) ATP is nearer the Smc3p-E1155 catalytic glutamate. **(Right side)** Acetylated cohesin inhibits Scc2p stimulation of cohesin ATPase: (1) Acetylation of Smc3p-K113 disrupts Scc2p-E822 binding, (2) leading to improper binding of Scc2p at the Smc1p-T1117 interface, (3) resulting in a failure to reposition the ATP closer to the catalytic glutamate, (4) thereby inhibiting Scc2p’s stimulation of cohesin’s ATPase.

These structural features of T1117, coupled with the functional features of the isoleucine suppressor substitution led us to hypothesize the following model for Scc2p’s activation of cohesin’s ATPase and its regulation by Smc3p-K113 acetylation. Without Scc2p binding to cohesin, the binding of ATP in the Smc3p ATPase active site was not optimal for ATP hydrolysis (Figure 4B). The binding of Scc2p at Smc3p-K113 helped orient Scc2p’s binding to Smc1p residues K1122 and N1096. The binding of Scc2p at this Smc1p interface repositioned the neighboring Smc1p-K1121 and ATP, such that ATP was more optimally positioned for hydrolysis by the catalytic glutamate in the Smc3 ATPase active site (Figure 4C, left side). Acetylation (or the acetyl mimic substitution) of Smc3p-K113 blocked its ability to form a salt bridge with Scc2p-E821 residue, thereby altering Scc2p binding at this interface and consequently Scc2p’s binding at the Smc1p-Scc2p interface (Figure 4C, right side). The altered Scc2p binding to Smc1p failed to induce the repositioning of Smc1p residues for optimal ATP binding and hydrolysis. The isoleucine substitution for T1117 acted as a surrogate for proper Scc2p binding, repositioning the Smc1-K1121 residue and the associated ATP to improve ATPase activity. Our model for Scc2p’s stimulation of cohesin’s ATPase and its regulation by Smc3p-K113 acetylation led us to four predictions about the in vivo and in vitro consequences of Smc1p-T1117 substitutions.

### Only smc1p-T1117I and smc1p-T1117V mutants suppress the inviability of the acetyl mimic

Our model predicted that only a few substitutions of the Smc1p-T1117 residue would promote the necessary subtle alteration of its neighboring residues to increase ATP hydrolysis and suppress the acetyl-mimic mutant defects. Most substitutions at T1117 would fail to suppress the defects of the smc3-K113Q mutant either because they failed to change the K1121 position, leading to no change in the low ATPase activity, or radically changed the K1121 position, leading to even worse ATPase activity. To test this prediction, we made a library of DNA repair templates that encoded all possible 19 amino acid substitutions at T1117 (Figure 5A). We assayed the ability of each of these substitutions to support the viability of otherwise wild-type cells (Supplemental Fig. 5A) and to suppress the inviability of the smc3-K113Q mutant (Figure 5B).

**Figure 5:**
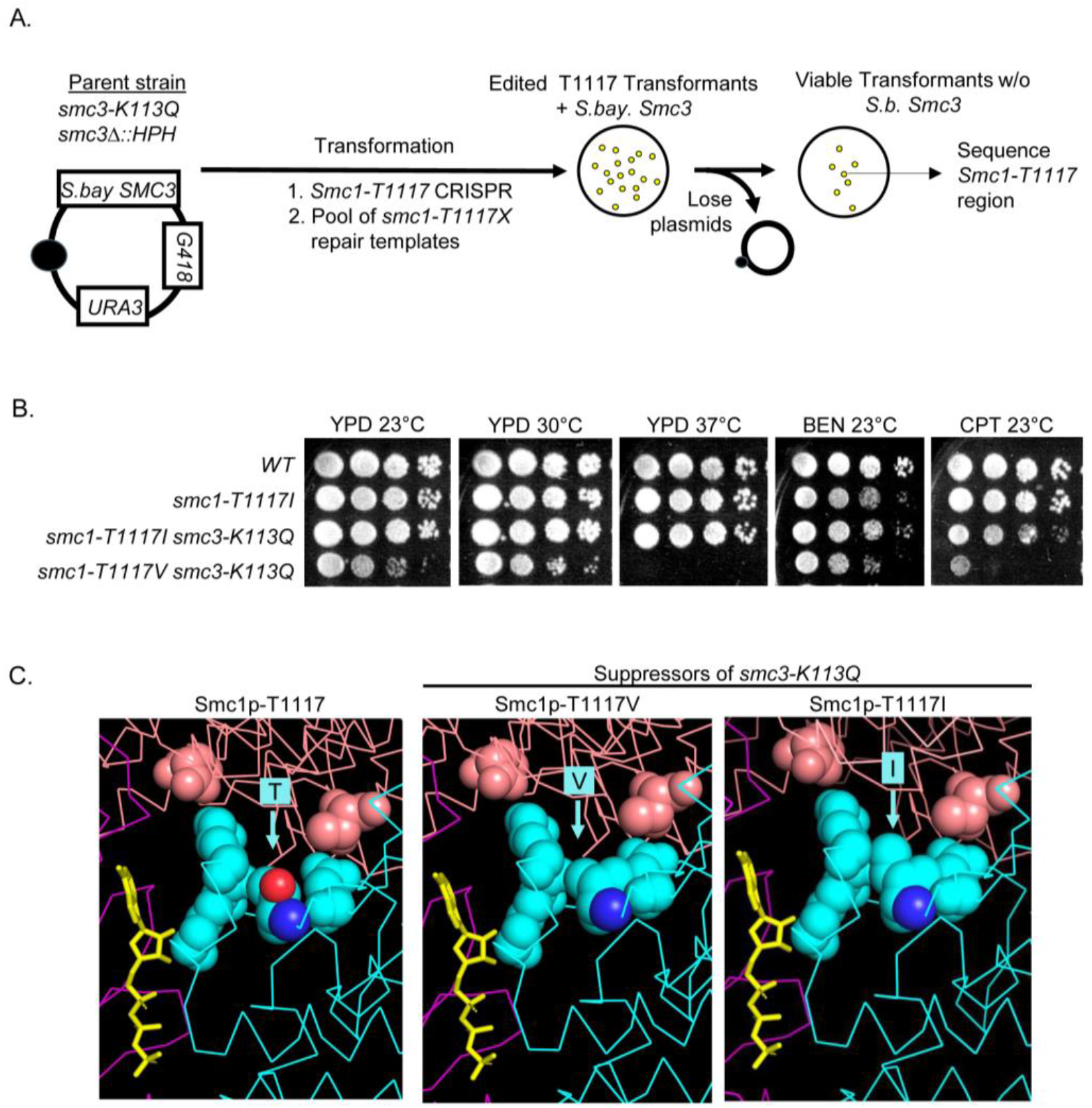
Only isoleucine or valine residue substitutions at smc1-T1117 suppress smc3-K113Q. **(A)** Schematic of a screen to assess which substitutions at smc1-T1117 suppress smc3-K113Q. Haploid strain (VG3969-14C) contains smc3-K113Q as the sole S. cerevisiae SMC3 and S. Bayanus SMC3 on a CEN URA3 G418 plasmid (pFC3). CRISPR was used to insert random substitutions of the smc1-T1117 residues as described in Materials and Methods. Transformants were replica plated to FOA media to select for loss of pFC3 (FOAR G418S colonies), which were sequenced to identify smc1-T1117 substitutions that suppresses smc3-K113Q inviability. **(B)** The valine substitution at smc1-T1117 (T1117V) is a weaker suppressor of smc3-K113Q than isoleucine (T1117I). Wild-type (VG3620-4C), smc1-T1117I (VG4006-13A), smc3-K113Q smc1-T1117I (VG4010-8B) and smc3-K113Q smc1-T1117V (VG4147-14C) were grown and diluted as described in Figure 1C then plated on YPD and incubated 4d at 23°C or 3d at 30°C and 37°C or on YPD containing benomyl [BEN] (10mg/ml) or camptothecin [CPT] (15mg/ml) and incubated for 4d at 23°C. **(C)** Cryo-EM of the Smc1p (cyan) T1117 interface with Scc2p (salmon). Wild-type Smc1p (left) and Smc3p-K113Q suppressor substitutions Smc1p-T1117V (middle), and Smc1p-T1117I (right).

In the wild-type background, all substitutions except proline were viable and grew well at 23°C and 37°C (Supplemental Fig. 5). However, tyrosine, glutamate, and aspartate substitutions were sensitive to benomyl and tyrosine to camptothecin, indicating they partially compromised cohesin function. The remaining fifteen substitutions were indistinguishable from the wild type for both drugs. By these criteria, most substitutions at smc1-T1117 appeared to be compatible with generating enough ATPase activity to support most, if not all, cohesin’s in vivo functions. However, valine was the only substitution at T1117, besides isoleucine, that could suppress smc3-K113Q inviability, albeit weaker than T1117I (Figure 4B). The fact that only two structurally similar substitutions, isoleucine and valine, could suppress the acetyl mimic was consistent with a subtle structural change that would be needed to improve ATPase activity. Modeling of the valine and isoleucine substitutions in the cryo-EM structure revealed that these substitutions subtly filled in the space between the ATP binding K1121 and the residues contacting Scc2p (Figure 5C). Thus, they could make a subtle change in the K1121 position that could alter ATP binding to improve its hydrolysis. In summary, these results showed that the smc3-K113Q suppressors at T1117 likely imposed a rare gain of function, consistent with the severe functional constraint of having to improve ATPase activity.

### The smc1p-T1117I substitution exacerbates the growth defect of acetyl-defective mutants

Previous studies suggested that the inviability and cohesion defects of mutants with unacetylated cohesin resulted from the failure to downregulate Scc2p stimulation of cohesin’s ATPase activity (Çamdere et al. 2015; Elbatsh et al. 2016; Murayama and Uhlmann 2015). If so, the growth defects of the acetyl-defective mutants should be made worse by smc1p-T1117I because it would further enhance the toxic ATPase activity of the unacetylated cohesin. To test this prediction, we introduced smc1-T1117I mutation into strains containing one of two conditional alleles of ECO1, either the temperature-sensitive eco1-(ctf7-203) or the auxin-sensitive ECO1-AID (Figure 6A, Supplemental Fig. 6A). We then tested these double-mutant strains for their growth under conditions where Eco1p function was reduced, leading to the under-acetylation of Smc3p-K113. The smc1-T1117I mutant not only failed to suppress the growth defects of the eco1 temperature-sensitive allele or the auxin-sensitive allele but exacerbated them further (Figure 6A, Supplemental Fig. 6A). This exacerbation fit with our model that the suppressor mutation altered the T1117 region of Smc1p to increase ATPase activity. This putative increase suppressed defects caused by acetyl-mimic mutant’s inhibition of ATPase stimulation but was toxic to cells with hyper-active ATPase generated when Smc3p-K113 acetylation levels were reduced.

**Figure 6:**
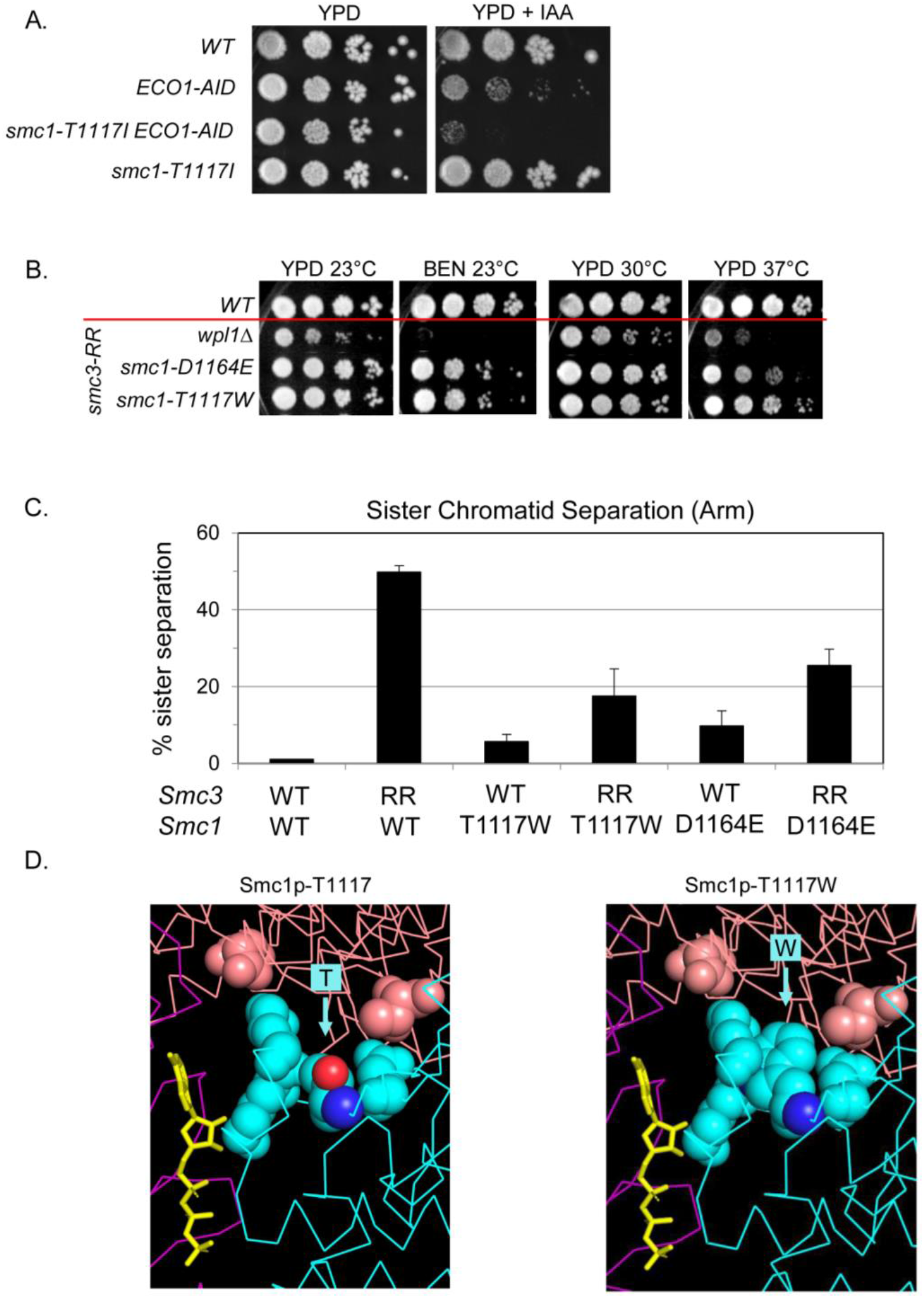
A tryptophan substitution at smc1-T1117 (T1117W) strongly suppresses the smc3-K112R, K113R (acetyl-null) mutant. **(A)** smc1-T1117I exacerbates the growth defect of ECO1-AID depletion. Wildtype (VG3620-4C), ECO1-AID (VG3633-3D), smc1-T1117I ECO1-AID (KB118E), and smc1-T1117I (JL11A) strains were grown and diluted as in Figure 1C then plated on YPD and YPD + IAA and incubated for 5d at 23°C. **(B)** smc1-T1117W and smc1-D1164E strongly suppress smc3-K112R, K113R (RR). Haploid wild-type (VG3620-4C), wpl1Δ smc3-K112R, K113R (VG4154-3A), smc1-D1164E smc3-K112R, K113R (VG4153-5C), smc1-T1117W smc3-K112R, K113R (VG4158-9D), were grown and diluted as in Figure 1C then plated on YPD and incubated at 23°C 4d, 30°C 3d or 37°C 3d and on YPD containing 10mg/ml benomyl and incubated at 23°C 5d (BEN 23°C). **(C)** smc1-T1117W strongly suppresses cohesion defect of the smc3-K112R, K113R mutant. Haploid wildtype (VG3620-4C), smc3-AID smc3-K112R, K113R (VG3991-1A), smc1-T1117W (VG4168-7B), smc1-T1117W smc3-K112R, K113R (VG4158-9D), smc1-D1164E (VG4138-1A), smc1-D1164E smc3-K112R, K113R (VG4153-5C) strains were grown as described in Figure 3B and processed to assess cohesion loss as described in Figure 1D. **(D)** Cryo-EM of Smc1p-T1117 (left) and acetyl-defective suppressor Smc1p-T1117W substitution (right).

### Substitutions of Smc1p-T1117 can also downregulate cohesin’s ATPase, acting as a surrogate for Smc3p acetylation

We reasoned that if the T1117 region of Smc1p was critical for mediating Scc2p’s stimulation of the ATPase, then a different subset of substitutions at T1117 might alter this critical region to decrease cohesin’s ATPase activity. These substitutions should behave similarly to Smc1p-D1164E, which also reduced cohesin ATPase activity and suppressed the in vivo defects of acetyl-defective mutants (Çamdere et al. 2015; Elbatsh et al. 2016).

To test this prediction, we asked whether any T1117 substitution could restore the biological and molecular functions of cohesin containing smc3p-K112R, K113R (Supplemental Fig. 6B). This acetyl defective mutant was known to be inviable and have a severe cohesion defect (Unal et al. 2008; Guacci et al. 2015). We identified eight substitutions at T1117 that suppressed the inviability of smc3-K112R, K113R (Supplemental Fig. 6C). Six of the eight substitutions (Ala, Asp, Glu, Gly, Ser, Lys) failed to grow at 37°C. They were extremely sensitive to benomyl (Supplemental Fig. 6B). The partial suppression of the growth defects of the acetyl-defective mutants by these substitutions was consistent with their reducing the ATPase activity of unacetylated cohesin sufficiently to support slow growth but not to restore all of cohesin’s biological functions.

In contrast, mutations to tryptophan (Trp) smc1-T1117W and phenylalanine (Phe) smc1-T1117F in the smc3-K112R, K113R background were nearly wild-type in their growth at high temperatures, and they also showed significant resistance to benomyl (Supplemental Fig. 6C). We compared the strongest acetyl-defective suppressor, smc1-T1117W to that of previously described acetyl-defective suppressors, wpl1Δ and smc1-D1164E. The smc1-T1117W suppressed the growth defects, benomyl sensitivity and cohesion defects of smc3-K112R, K113R much better than the wpl1Δ and slightly better than smc1-D1164E (Figure 6B-C, (Çamdere et al. 2015)). Modeling of these tryptophan and phenylalanine substitutions at T1117 revealed that they clashed with the position of K1121 (Figure 6D) providing a mechanism for how they could alter the positioning of the ATP in the active site and reduce its hydrolysis.

Taken together, our in vivo suppressor analyses showed that different subsets of T1117 substitutions could either enhance cohesin function to counter its downregulation by acetylation, or reduce cohesin function to counter its constitutive upregulation by lack of acetylation. These results were consistent with our model that the T1117 region of Smc1p was capable of toggling cohesin’s ATPase levels in response to proximal Scc2p binding and the acetylation state of Smc3p-K113 residue.

### smc1p-T1117I increases cohesin ATPase activity

Our model also predicted that cohesin’s ATPase activity in vitro should be enhanced by Smc1p-T1117I. To test this prediction, we purified wild-type and mutant cohesins from an eco1Δ wpl1Δ strain that was alive but unable to acetylate cohesin. We measured their ATPase activity in the presence of DNA without Scc2p (basal) and with Scc2p (induced). We observed a similar basal ATPase activity for cohesin with wild-type Smc3p or Smc3p-K113Q (Figure 7A-B). Scc2p addition stimulated ATPase activity of WT cohesin 4-5-fold but failed to stimulate Smc3p-K113Q cohesin as reported previously (Figure 7A, (Murayama and Uhlmann 2015)). The presence of Smc1p-T1117I increased cohesin’s basal ATPase activity about two-fold (Figure 7A). Addition of Scc2p increased ATPase activity of smc1p-T1117I cohesin to a level 50% greater than wild-type cohesin (Figure 7A). For cohesin containing both Smc3p-K113Q and Smc1p-T1117I, the basal ATPase activity was dramatically increased relative to cohesin with just Smc3p-K113Q (Figure 7A). This double mutant cohesin reached ATPase levels greater than that seen for wild-type cohesin with Scc2p (Figure 7A). Thus, as predicted from our model, the smc1-T1117I substitution restored cohesin ATPase activity to cohesin with the acetyl-mimic mutation.

**Figure 7:**
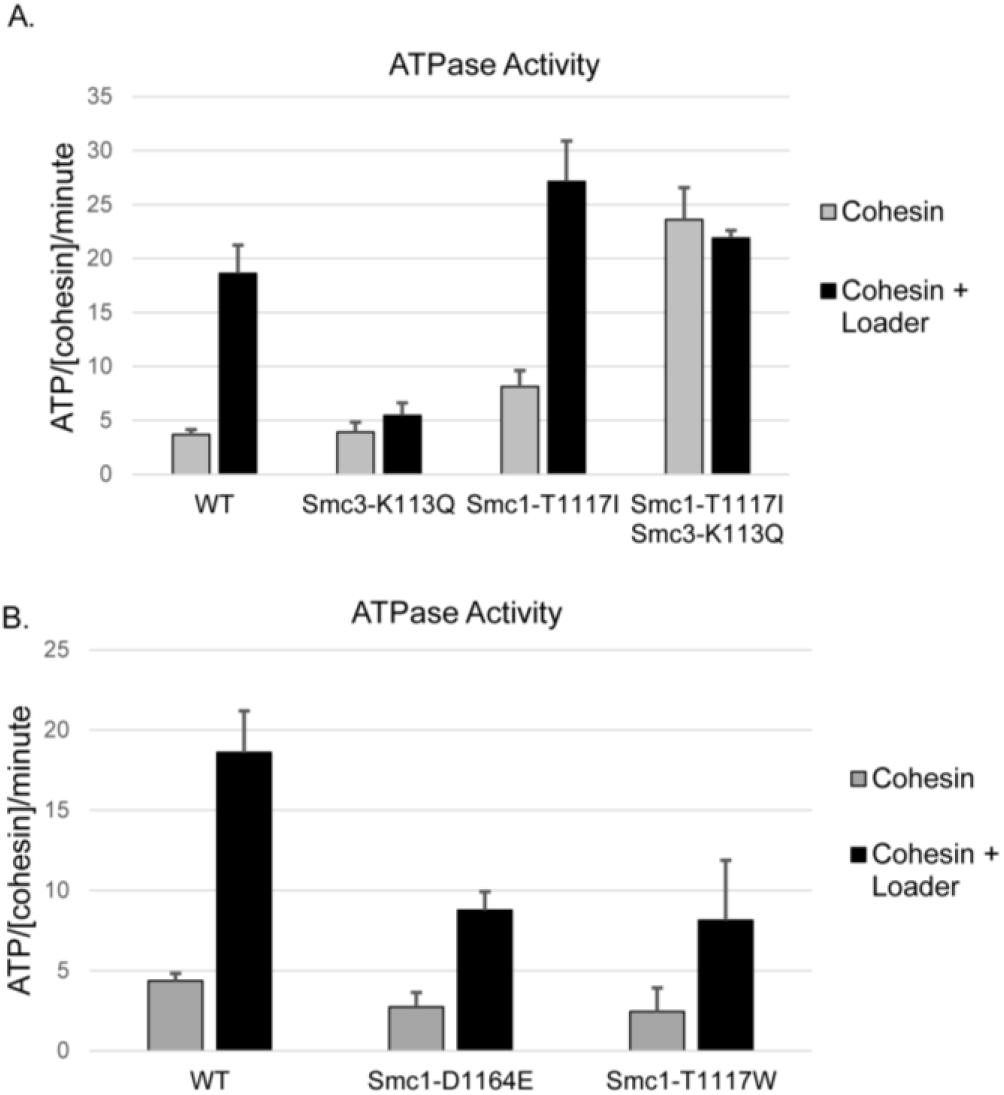
Different substitutions at SMC1-T1117 up-regulate or down-regulate cohesin ATPase activity. **(A)** The Smc1p-T1117I suppressor of the Smc3p acetyl-mimic up-regulates cohesin ATPase activity. Wild-type and mutant cohesin complexes were purified and assessed for ATPase activity with and without Scc2p/Scc4p (Loader). Purified cohesin concentration normalization was confirmed (Supplemental Fig. 7A). **(B)** The Smc1p-T1117W and Smc1p-D1164E suppressors of the Smc3 acetyl-null down-regulate cohesin ATPase activity. Wild-type and mutant cohesin complexes were purified and assessed for ATPase activity with and without Scc2p/Scc4p (Loader). Purified cohesin concentration normalization was confirmed (Supplemental Fig. 7B).

As a corollary, our model also predicted that cohesin’s ATPase should be inhibited by Smc1p-T1117W similar to Smc1p-D1164E. Cohesin with these two Smc1p mutants had similar levels of basal cohesin ATPase. Scc2p addition induced ATPase activity of both mutant complexes 2-fold less than wild-type, similar to the levels previously reported for cohesin with Smc1p-D1164E (Elbatsh et al. 2016). Therefore, Smc1p-T1117W reduced cohesin ATPase activity as our in vivo results and model predicted.

An unexpected observation from these studies was that the Scc2p-independent ATPase activity of cohesin with both Smc3p-K113Q and Smc1p-T1117I was significantly greater than cohesin with Smc1p-T1117I alone or Smc3p-K113Q alone. Thus, this very high level of basal ATPase activity required Smc3p-K113Q as well as the Smc1p-T1117I substitution. This result suggested that the acetylation state of Smc3p-K113 could potentially have additional impact on cohesin’s ATPase activity beyond modulating Scc2p function.

The increase in Scc2p-independent ATPase activity of cohesin by Smc1p-T1117I also suggested that this substitution might reduce, or perhaps even bypass, the need for Scc2p in vivo. To test this possibility, we introduced the smc1-T1117I allele into cells harboring either the auxin-sensitive SCC2-AID or temperature-sensitive scc2-4 alleles. The growth on auxin (IAA)-containing media of the SCC2-AID smc3-T1117I mutant was significantly greater than the SCC2-AID mutant but not restored to wild-type growth (Supplemental Fig. 7C). The temperature-sensitive growth of the scc2-4 was also partially suppressed by smc1-T1117I as evidenced by growth at 30°C but not 35°C (Supplemental Fig. 7D). However, the SCC2-AID smc1-T1117I double mutant was identical to SCC2-AID alone as both strains exhibited extremely poor cohesion and very low cohesin binding to chromosomes (Supplemental Fig. 7E-G). Finally, the smc1-T1117I smc3-K113Q double mutant was unable to suppress a loss of Scc2p (Supplemental Fig. 7H). Together, these results suggested that higher basal ATPase of smc1-T1117I cohesin partially compensated for reduced Scc2p function but even the hyper-active smc3-K113Q smc1-T1117I cohesin cannot bypass the need for Scc2p.

## Discussion

In this study, we sought to answer three mechanistic questions about cohesin regulation. How does Scc2p activate cohesin’s ATPase? Does acetylating Smc3p-K113 impede the ability of Scc2p to activate cohesin ATPase in vivo, as suggested by in vitro experiments? How is cohesin’s ATPase activity repressed by Smc3p-K113 acetylation given that this residue is not proximal to either of cohesin’s ATPase active sites? To answer these questions, we isolated suppressors of the inviability of smc3-K113 acetyl-mimic or acetyl-null mutants. We assayed the impact of these mutant proteins on cohesin function through in vitro ATPase assays, and in vivo assays of cohesion, chromosome-binding, drug resistance, and cohesin structural integrity. We also interrogated how suppressors impacted cohesin structure using the cryo-EM structure of cohesin with Scc2p. From our results, we developed a model for Scc2p stimulation of cohesin ATPase and its regulation by acetylation of the Smc3-K113 residue. Below, we summarize the model and our key observations that support the model.

We propose that Scc2p’s interaction with Smc3p at its K113 residue affects the interface of Scc2p with Smc1p residues that lie near the Smc3p ATPase active site. The binding at this Scc2p-Smc1p interface causes a shift in the neighboring Smc1p residues and the nearby ATP bound in the Smc3p ATPase active site such that the ATP is better oriented for hydrolysis by the catalytic glutamate of Smc3p. Acetylation of Smc3p-K113 directly alters Scc2p’s interface with Smc3p, which in turn alters Scc2p’s interface with Smc1p residues proximal to the Smc3p ATPase site. These acetylation-mediated alterations inhibit Scc2p’s ability to shift the key residues in Smc1p that promote ATP hydrolysis.

This model is based on four key observations. First, we showed that overexpression of the Scc2p-Scc4p complex weakly suppressed the inviability of the smc3-K113Q mutant (acetyl-mimic). This result indicates that the acetylation of cohesin limits Scc2p function in vivo as had been suggested by a Cryo-EM structure and in vitro studies of Scc2p activation of cohesin ATPase (Murayama and Uhlmann 2014; Shi et al. 2020). This partial suppression by Scc2p overexpression also suggests that Smc3p-K113 acetylation does not abolish Scc2p binding to the cohesin head, but rather alters the interaction. Second, all the suppressors of the acetyl-mimic lie in the Smc head domains proximal to the Smc3p ATPase active site, suggesting that Smc3p acetylation regulates cohesin specifically by modulating the Smc3p ATPase activity. Similarly, previously identified suppressors of Smc3p acetyl defective mutants also lie proximal to the Smc3p ATPase (Çamdere et al. 2015; Elbatsh et al. 2016). Third, we demonstrate that substitutions of the Smc1p-T1117 residue were capable of toggling cohesin’s ATPase levels up or down, suppressing Smc3p acetyl-mimic or acetyl-defective mutants, respectively. This residue is proximal to Smc1p residues that interface with Scc2p and that help position ATP in the Smc3p ATPase active site. Structural modeling of these substitutions in the cryo-EM structure reveals that they would cause subtle changes that alter ATP positioning in the Smc3p ATPase active site, consistent with their impact on ATPase activity. Thus, it is reasonable to propose that Scc2p binding proximal at this Smc1p region could induce subtle structural changes similar to those induced by the acetyl-mimic suppressors to enhance ATPase activity. Finally, different substitutions at smc1-T1117 suppress the inviability of the acetyl-mimic and defective mutants of Smc3p-K113. This fact connects acetylation-dependent alterations in Scc2p’s interaction with the Smc3p-K113 residue to the distal interactions of Scc2p with the critical Smc1p region that controls ATPase activity.

Our model raises a conundrum. Why not control the ATPase by modifying the Scc2p-Smc1p interface, which directly modulates ATPase activity rather than by modifying a distal interface between Scc2p and the Smc3p-K113 residue? One possibility is that differentially regulating cohesin’s loop extrusion, stable loop formation, and cohesion functions require communication between biochemical activities in the Smc3p-Scc2p region to the Smc3p ATPase. Indeed, a recent study suggests a model for loop extrusion (Bauer et al. 2021), wherein DNA is entrapped between Scc2p and the cohesin head near the Smc3p-K113-Scc2p and also binds at the hinge. Transfer of DNA from the hinge to the head is thought to be dependent on Scc2p’s stimulation of cohesin’s ATPase. Cycles of ATP binding and hydrolysis would allow the cycles of DNA release and recapture that are needed for loop extrusion. Inhibiting Scc2p’s stimulation of cohesin’s ATPase by Smc3p-K113 acetylation would trap DNA binding to the hinge, generate a stable DNA loop from loop extrusion, or generate cohesion if the hinge is bound to the sister chromatid. Thus, sensing DNA binding in the Smc3p-K113 region could be an important input to stimulate ATPase for loop extrusion and the acetylation-dependent entrapment of DNA needed for tethering.

Another conundrum arises from our suppressor analysis. In wild-type cells, only a fraction of cohesin is acetylated, presumably to generate two pools of cohesin, one with high ATPase for looping and another with low ATPase for cohesion. However, the smc1-T1117I and mc1-T1117W mutations allow near perfect suppression of the defects caused by acetyl-mimic and null mutations respectively. These results suggest that all cohesin’s biological functions can be carried out by cohesin in a single acetylated state with ATPase activity either lower (smc3-K112R, K113R smc1-T1117W, or smc3-K113R smc1-D1164E) or higher (smc1-T1117I smc3-K113Q) that wild type. One possibility is that the ATPase and its biological functions are controlled by an additional regulatory protein like Pds5p (Bastié et al. 2022; van Ruiten et al. 2022). However, invoking redundancy avoids the question of what is the fitness advantage for the acetylation-dependent control of cohesin’s ATPase. A clue may come from the one residual phenotype in these cells with fixed ATPase levels: they exhibit some sensitivity to DNA-damaging agents. It will be exciting to use the different suppressors to probe the impact of the different acetylation and ATPase states on DNA repair and chromosome structure.

## Supporting information

Supplemental Table 1 - Strain List

Supplemental Table 2 - Plasmid List

Supplemental Table 3 - Primer List

## Acknowledgements

We thank Jasper Rine and Lorenzo Costantino for critical reading of the manuscript. We thank Jonathan Luo for help with strain construction. This work was funded by the National Institutes of Health Grant (1R35 GM-118189-06 to DK).

## Author contributions

K.M.B characterized the Smc3p acetyl-mimic and the effect that Scc2p/Scc4p overexpression had on the acetyl-mimic. V.G. & F.C conducted the screen for suppressors of the acetyl-mimic mutant and the preliminary characterization of all the suppressors. K.M.B. characterized in detail the smc1-T1117I mutant alone, its suppression of the acetyl-mimic, and its effect on eco1 mutants. V.G. assessed whether smc1-T1117I bypasses the need for Scc2p. K.M.B. produced sequence alignments. K.M.B and D.K produced molecular modeling on the S. cerevisiae and H. sapiens cryo-EM structures. D.K. produced the cartoon model. V.G. isolated and characterized substitutions at smc1-T1117 that suppress an Smc3p acetyl-null or acetyl-mimic. U.M. isolated and characterized substitutions at smc1-T1117 that support viability in wild-type cells. K.M.B. assisted S.X. with construction of cohesin purification strains and purification of wild-type cohesin. S.X. purified mutant cohesin complexes. S.X. assayed ATPase activity of wild-type and mutant cohesin complexes. D.K. was the main writer of the manuscript. K.M.B. and V.G. were major editors of the manuscript prose and organization and preparation of the figures.

## Materials and Methods

### Yeast strains, media, and reagents

Yeast strains used in this study are A364A background, unless otherwise specified. Genotypes are listed in Supplemental_Table_1. YPD media was made as previously described (Guacci et al. 1997). YEPR or YEPG are the same as YPD except contain 2% raffinose or galactose instead of dextrose. YEPRG has 2% galactose and raffinose. Plates containing benomyl or camptothecin (Sigma catalog # C9911) used to assess drug sensitivity were prepared as previously described (Guacci and Koshland 2012). Auxin (3-indoleacetic acid) (Sigma-Aldrich, St. Louis, MO) was made as a 1M stock in DMSO then added to 500μM or 750μM final concentration in liquid media or plates, respectively.

#### Cohesin Purification Media

Low Biotin Synthetic Complete (LBSC) Media contained 1.56 g/L BSM Powder (Sunrise Science Products Cat#1387), 1.71 g/L YNB – Biotin powder (Sunrise Science Products Cat#1523), 38 mM ammonium sulfate (5 g/L), 1 nM D-biotin (Invitrogen #B20656) and 2% raffinose.

#### Cohesin Loader Purification Media

Low Biotin URA-Dropout Media contained 0.8g/L CSM-Ura (Sunrise Sience Products), 1.71g/L YNB –Biotin powder (Sunrise Science Products Cat#1523), 38 mM ammonium sulfate (5g/L), 1 nM biotin and 2% raffinose.

### Dilution plating assays

Cells were grown to saturation in YPD media at 23°C or 30°C, plated in 10-fold serial dilutions on YPD alone or containing drugs then incubated at 23°C or 30°C.

### G1 arrest and synchronous release into mid-M phase arrest

#### G1 arrest

Asynchronous mid-log cultures were arrested in G1 by addition of alpha factor as previously described (Guacci et al 2019). When required, auxin was added (500mM final) to G1 arrested cells, incubated 30 minutes while arrested in G1. To induce pGAL promoters, galactose was added to 2% final and cells incubated 30 minutes.

#### Synchronous release from G1 into mid-M phase arrest

G1 arrested cells were released from G1 into either YPD or YEPRG containing nocodazole and Pronase E as previously described (Guacci et al. 2019) then incubated at 30°C for 2.5h for YPD or 4h for YEPRG to arrest in mid-M phase. When required, auxin was added (500mM final) in all wash media and in resuspension media to ensure AID-tagged protein depletion.

### Protein extracts and western blotting

### Total protein extracts

2 to 4 OD600 cells equivalents were frozen and protein extracts made as described in (Guacci et al. 2019).

### Western blots

Protein extracts were loaded on 8% SDS-page gels, subjected to electrophoresis then transferred to PDVF membranes. Proteins were detected using HRP-conjugated antibodies.

### Chromatin immunoprecipitation (ChIP)

Aliquots of cells synchronously arrested in mid-M phase were fixed and processed for ChIP as described previously (Guacci et al. 2019). Primers used for ChIP are shown in Supplementary_Table_3.

### Monitoring cohesion using LacO-GFP assay

Cohesion was monitored at CEN-proximal and distal loci using the LacO-LacI system as previously described (Guacci and Koshland 2012). Mid-M phase cells were fixed, and the number of GFP signals in each cell scored. Cells with 2 GFP spots have defective cohesion. G1-arrested cells were scored to confirm that mid-M phase cells with 2 GFP spots were not due to pre-existing aneuploidy.

### Microscopy

All images were collected on a Zeiss Axio Observer Z1 microscope equipped with Plan-Apochromat 63x objective and the Definite Focus System to maintain focus over time. z-series optical sections were collected with a step-size of 0.2 microns, using an ASI MS-2000 XYZ piezo stage. Multiple stage positions were collected. The microscope, camera, and stage were controlled with the Micro-manager software (Edelstein et al. 2014). Z-series were viewed using Fiji software (Schindelin et al. 2012).

### Flow cytometry

Flow cytometry analysis was performed as previously described (Bloom et al., 2018), except that the fixed cells were washed twice in 1X TE with 0.2% Tween-20 (v/v).

### CRISPR-mediated strain building

CRISPR guide plasmids and PCR generated repair templates were made and used to insert mutations into yeast as previously described (Saxton and Rine 2019). CRISPR guides and repair templates are listed in Supplementary_Table_2 and Supplementary_Table_3.

### Isolation of spontaneous suppressors of the smc3-K113Q mutant

#### Screen for suppressors

Haploid VG3969-14C contains pFC3 (S. bayanus SMC3 CEN URA3 G418), endogenous SMC3 deleted (smc3Δ::HPH), and smc3-K113Q integrated at LEU2 (pVG419 K113Q). S. Bayanus SMC3 gene supports viability and prevents gene conversion of smc3-K113Q. Single colonies were grown to saturation in YPD, washed with H2O, plated 5-FOA (FOA) and incubated at 30°C. FOAR G418S colonies have lost pFC3. PCR sequencing confirmed smc3-K113Q remained, so colonies have suppressor mutations. These were sequenced to identify the suppressor.

### Rebuild strains to confirm suppressors

#### SMC3 suppressors

Haploid 3961-4B (smc3Δ::HPH + pFC3 [S.bay. SMC3 CEN URA3 G418]) was transformed with PpuMI-linearized LEU2 plasmids containing WT SMC3 (pVG419), a suppressor allele alone or suppressor with smc3-K113Q. pFC3 was lost by plating strains on 5-FOA media. Mutants were confirmed by PCR sequencing.

#### SMC1 suppressors

Haploid 3965-1A (smc1Δ::HPH + pFC1 [S.bay. SMC1 CEN URA3 G418]) was transformed with PpuMI-linearized LEU2 plasmids containing WT SMC1 (pVG444) or bearing the suppressor smc1 alleles. CRISPR was used to insert smc3-K113Q at the endogenous locus in strains then pFC1 was lost by plating on 5-FOA media. Mutants were confirmed by PCR sequencing.

### Screen for smc1-T1117 residues that suppress smc3-K113Q

CRISPR was used to insert random residues at smc1-T1117 (PCR repair smc1-T1117X) in haploid 3961-4B (smc3-K113Q-LEU2:leu2-3,112 smc3Δ::HPH + pFC3 [S.bay. SMC3 CEN URA3 G418]). Transformants were plated on media to select for CRISPR plasmid and colonies replica plated to FOA media. FOAR colonies have lost pFC3 so contain smc1-T1117 substitutions that suppress smc3-K113Q. PCR sequencing confirmed smc3-K113Q remained and identified which smc1-T1117 residue substitutions are suppressors. Thirteen strong (drug resistant) and 9 weak (drug sensitive, temperature sensitive) suppressors were sequenced. All strong suppressors were isoleucine (T1117I) and all weak suppressors were valine (T1117V).

### Assessing which substitutions at smc1-T1117 support viability in WT cells

CRISPR was used to insert random residues at smc1-T1117 (smc1-T1117X) into wild-type haploid 3620-4C. PCR sequencing of transformant colonies identified smc1-T1117 residues that support viability.

### Screen for smc1-T1117 residues that suppress smc3-K112R, K113R

CRISPR was used to insert random residues at smc1-T1117 (smc1-T1117X) into haploid 4144-5C (smc3-K112R,K113R-LEU2:leu2-3,112, smc3Δ::HPH + pFC3 [S.bay. SMC3 CEN URA3 G418]). Smc1-T1117 residues that suppress smc3-K112R, K113R were identified as described above to identify for smc1-T1117 residues that suppress K113Q.

### Sequence Alignment

Protein amino acid sequences were aligned in AliView using MUltiple Sequence Comparison by Log-Expectation (MUSCLE).

### Protein Purification

#### Cohesin Purification

Cohesin purification strains were grown in Low Biotin Synthetic Complete Media containing 2% raffinose to OD 1.0 at 30 °C. Galactose was added to 2% to induce protein expression and incubated at 30°C for 2h. Cells were collected by centrifugation, washed once with cold PBS, washed with lysis buffer (50mM HEPES pH 7.5, 10% (v/v) glycerol, 150mM NaCl, 5mM MgCl2, 0.1mM CaCl2), pelleted then frozen. The frozen pellet was thawed on ice and resuspended in lysis buffer containing 1.2% IGEPAL CA-630, 20mM β-Mercaptoethanol, and EDTA-free protease inhibitor tablet (Sigma). DNase I (Millipore-Sigma #11284932001) was added to a final concentration of 0.05 mg/ml, then PMSF added to a final concentration of 0.25mM. Cells were lysed by sonication then lysate clarified by centrifugation at 20,000x g for 45min at 4°C. Clarified lysate was loaded onto a StrepTrap XT column (Cytiva), pre-equilibrated with lysis buffer. The column was washed with 10 column volumes of lysis buffer then eluted with 6 column volumes of Elution buffer 1 (50mM HEPES pH7.5, 10% (v/v) glycerol, 2mM MgCl2, 300mM NaCl, 50mM D-biotin, 20mM β-Mercaptoethanol). Eluate was loaded onto a HiTrap Heparin HP column (Cytiva) then eluted using 5 column volumes of Elution Buffer 2 (50mM HEPES pH7.5, 10% (v/v) glycerol, 2mM MgCl2, 800mM NaCl, 20mM β-Mercaptoethanol), only the last 4 column volumes were collected. NaCl concentration was adjusted to 266mM NaCl then protein concentrated by ultrafiltration (Thermo Scientific Pierce Protein Concentrater PES 30K Cat# PI88529S).

#### Cohesin Loader Purification

The loader purification strain was grown in Low Biotin URA-Dropout Media containing 2% raffinose to OD 1.0 at 30°C. Galactose was added to 2% to induce protein expression and incubated at 30°C for 2h. Cells were collected by centrifugation, washed once with cold PBS, then washed with lysis buffer (50mM HEPES 7.5, 10% (v/v) glycerol, 150mM NaCl, 5mM MgCl2, 0.1mM CaCl2), then the pellet was frozen. The frozen pellet was thawed on ice and resuspended in lysis buffer containing 1.2% IGEPAL CA-630, 20mM β-Mercaptoethanol, and EDTA-free protease inhibitor tablet (Sigma). DNase I was added to a final concentration of 0.05 mg/ml, then PMSF added to a final concentration of 0.25 mM. Cells were lysed by sonication then lysate clarified by centrifugation at 20,000 g for 45min at 4°C. Clarified lysate was loaded onto a StrepTrap XT column, pre-equilibrated with lysis buffer. The column was washed with 10 column volumes of lysis buffer, followed by 15 column volumes of STW Buffer 300 (50mM HEPES 7.5, 10% (v/v) glycerol, 2mM MgCl2, 300mM NaCl) then eluted with 5 column volumes of LE Buffer 1 (50mM HEPES pH 7.5, 20% glycerol, 5mM MgCl2, 300mM NaCl, 50mM Biotin, 20mM β-Mercaptoethanol). Eluate was loaded onto a HiTrap Heparin HP column (GE Healthcare), washed with 6 column volumes of LHW Buffer 300 (50mM HEPES pH 7.5, 20% (v/v) glycerol, 2mM MgCl2, 300mM NaCl) then eluted using 5 column volumes of LE Buffer 2 (50mM HEPES 7.5, 20% (v/v) glycerol, 2mM MgCl2, 800mM NaCl, 20mM β-Mercaptoethanol). Only the last 4 column volumes were collected. NaCl concentration was adjusted to 266mM NaCl using LHW Buffer 0 (50mM HEPES 7.5, 20% (v/v) glycerol, 2mM MgCl2, 20mM β-Mercaptoethanol) and protein concentrated by ultrafiltration (Thermo Scientific Pierce Protein Concentrator PES 30K Cat# PI88529S).

#### ATPase Assay

ATPase activity of cohesin was measured using EnzChek Phosphate Assay Kit with purified recombinant proteins depleted of free phosphate using Inorganic Phosphate Binding Resin (Abcam: ab270547). Reactions were assembled with10nM cohesin, 15nM Scc3, alone or with 65nM Scc2/4, 0.1mg/ml BSA and 450nM 60-mer dsDNA in ATPase reaction buffer (25mM HEPES pH7.5, 20% glycerol, 50mM NaCl, 1mM MgCl2); reactions were initiated with addition of ATP to a final concentration of 1mM. Spectrophotometric measurements at 360nM were taken every 1min for 2h at room temperature. ATPase activities were calculated by linear regression of the raw data using GraphPad Prism software.

**Supplemental Figure 1.**
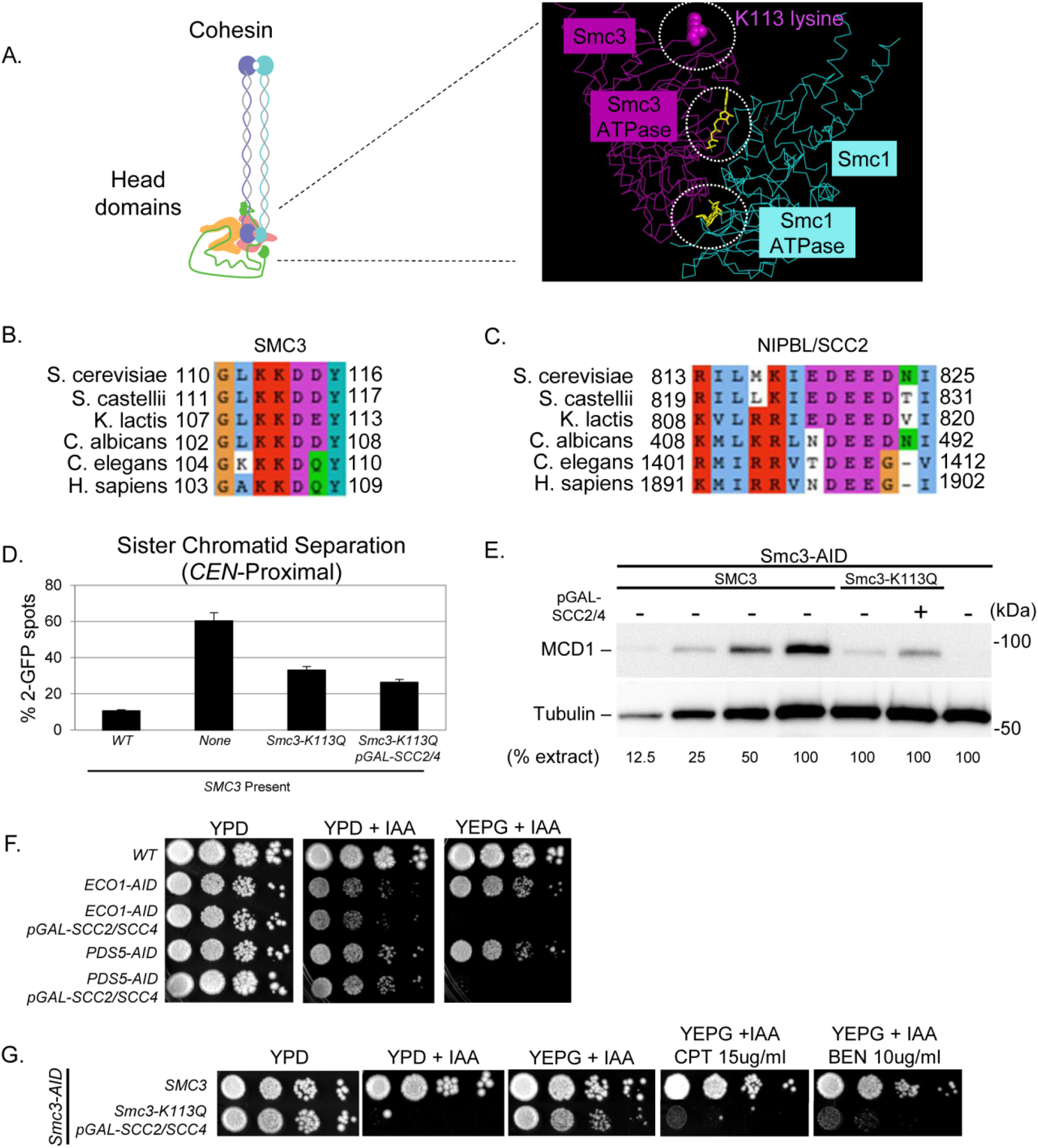
**(A)** Cartoon depiction of cohesin (left). S. cerevisiae cryo-EM of the head domains of Smc1p (cyan) and Smc3p (magenta) with ATPases, ATP (yellow) and the Smc3-K113 residue shown (right). **(B)** Amino acid sequence alignment of the S. cerevisiae Smc3p-K113 region shown in Figure 1B with other Smc3p orthologs including H. sapiens Smc3p. **(C)** Amino acid sequence alignment of the S. cerevisiae Scc2p-E822 region shown in Figure 1B with other Scc2p orthologs including H. sapiens NIPBL. **(D)** SCC2/SCC4 overexpression fails to suppress the CEN-proximal cohesion defect of smc3-K113Q cells. Haploid strains bearing SMC3-AID alone (JL16A) or also containing wild-type SMC3 (JL17A), smc3-K113Q (JL14A), or smc3-K113Q with pGAL-SCC2/SCC4 (JL15A) were grown and fixed as in Figure 1D. Strains contained LacO 10kb from CEN4 to assess cohesion at a CEN-proximal locus. Cohesion was assessed as described in Figure 1D. **(E)** Mcd1p levels are reduced when the smc3-K113Q acetyl-mimic mutant is the sole Smc3p present, but SCC2/SCC4 overexpression partially restores Mcd1p levels. Protein extracts (TCA lysed) from 2 OD of mid-M phase cells in Figure 1D and 1E were analyzed by Western blot. Mcd1p protein levels were monitored using rabbit antibodies against Mcd1p (αMCD1) and rabbit antibodies against tubulin (αTUB2) as a loading control. **(F)** SCC2/SCC4 overexpression does not suppress ECO1-AID or PDS5-AID growth defects. Haploid wild-type (KB61A), ECO1-AID alone (VG3633-2D), ECO1-AID containing pGAL-SCC2/SCC4 (KB50A), PDS5-AID alone (VG3862-1A), and PDS5-AID containing pGAL-SCC2/SCC4 (KB46A), were grown were grown and dilution plated as in Figure 1C on YPD, YPD + IAA, or YEPG + IAA and incubated for 4 days at 23°C. **(G)** SCC2/SCC4 overexpression partially suppresses the inviability of smc3-K113Q. Haploid SMC3-AID strain also bearing either a wild-type SMC3 (VG3919-3C), or smc3-K113Q containing pGAL-SCC2/SCC4 (VG4052-3A), were grown and diluted as in Figure 1C then plated on YPD, YPD + IAA, or YEPG+ IAA, YEPG + IAA + camptothecin [CPT] (15mg/ml), YEPG + IAA + benomyl [BEN] (10mg/ml) and incubated for 4 days at 23°C.

**Supplemental Figure 2.**
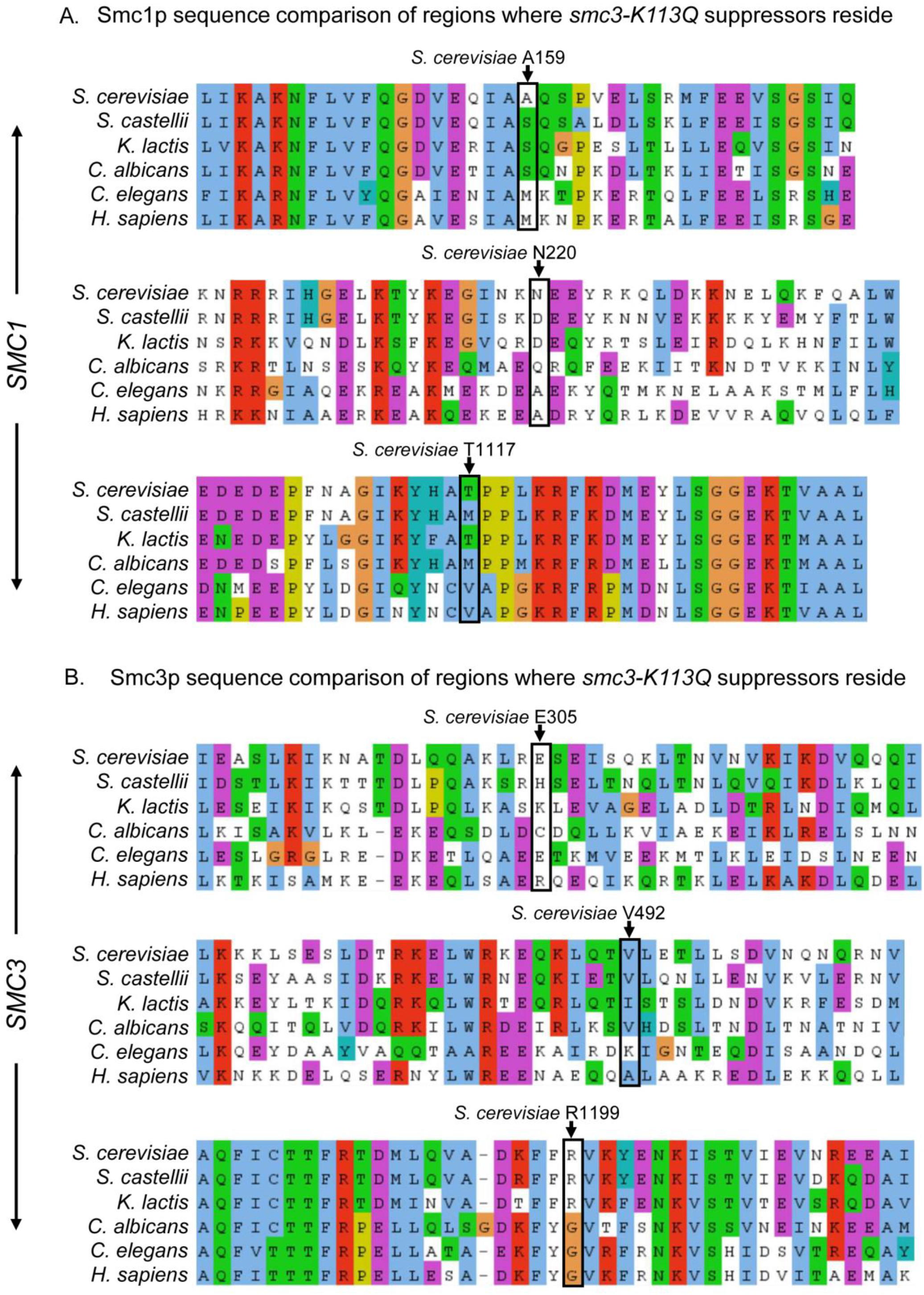
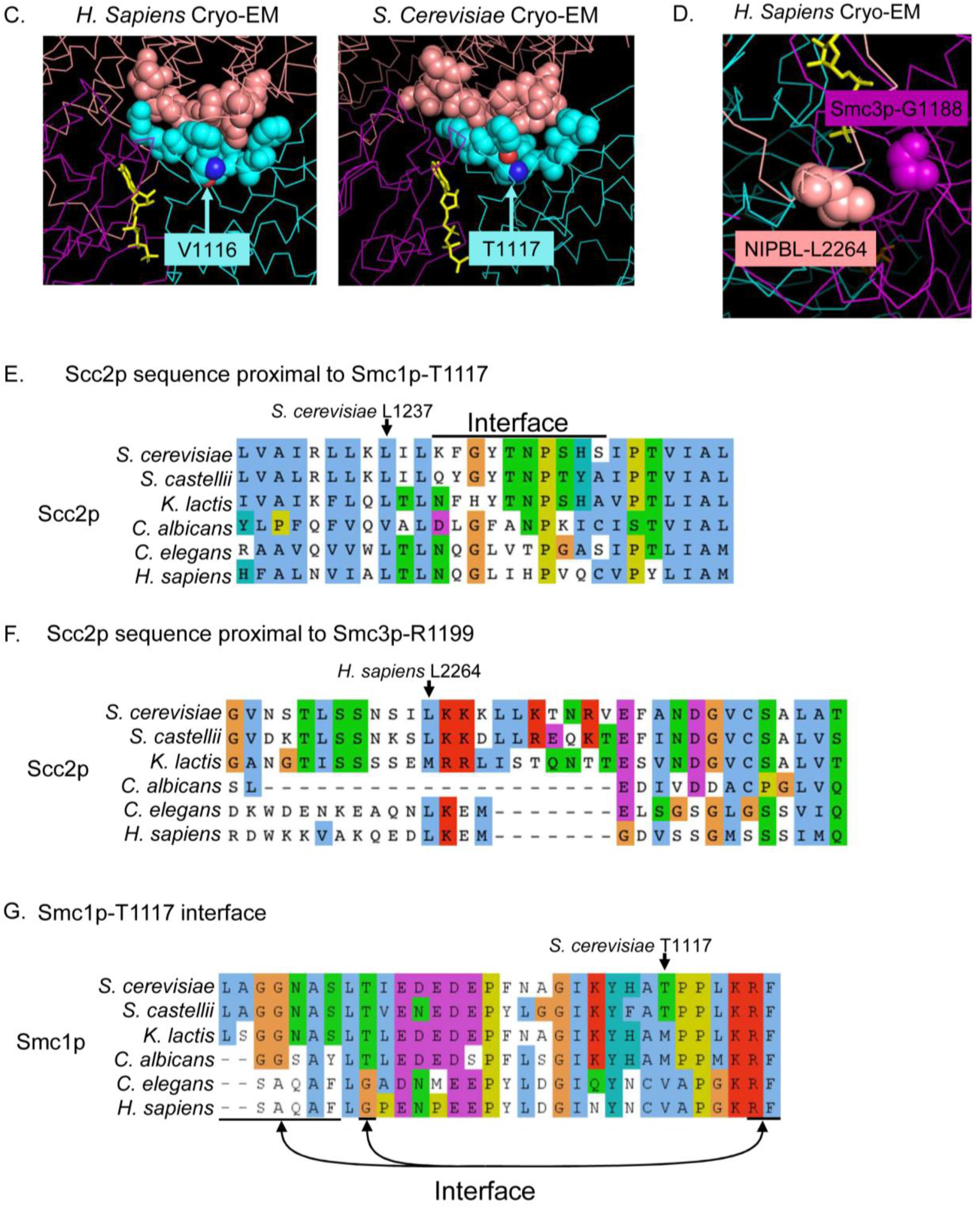
**(A)** Amino acid sequence alignment of Smc1p orthologs indicating the positions of the three suppressor S. cerevisiae residues: A159, N220 and T1117. **(B)** Amino acid sequence alignment of Smc3p orthologs indicating the positions of the three suppressor S. cerevisiae residues: E305, V492 and R1199. **(C)** Cryo-EM of H. sapiens (left) and S. cerevisiae (right) indicating Smc1p (cyan) and Scc2p (salmon), and ATP (yellow). Residues shown as spheres illustrate where the Smc1p-T1117 region interfaces with Scc2p. **(D)** H. sapiens cryo-EM of Smc3p-G1188 (magenta) interfaces with NIPBL (Salmon) indicated as spheres. shown as spheres illustrate where the Smc1p-T1117 region interfaces with Scc2p. ATP is yellow. **(E)** Amino acid sequence alignment of Scc2p orthologs showing the region which interfaces with S. cerevisiae Smc1p-T1117. **(F)** Amino acid sequence alignment of NIPBL/Scc2p orthologs showing the H. sapiens Scc2p region that interfaces with Smc3p-G1188 (Smc3p-R1199 in S. cerevisiae). **(G)** Amino acid sequence alignment of Smc1 orthologs showing the Smc1p-T1117 region. S. cerevisiae Smc1p residues which interfaces with Scc2p are indicated.

**Supplemental Figure 3.**
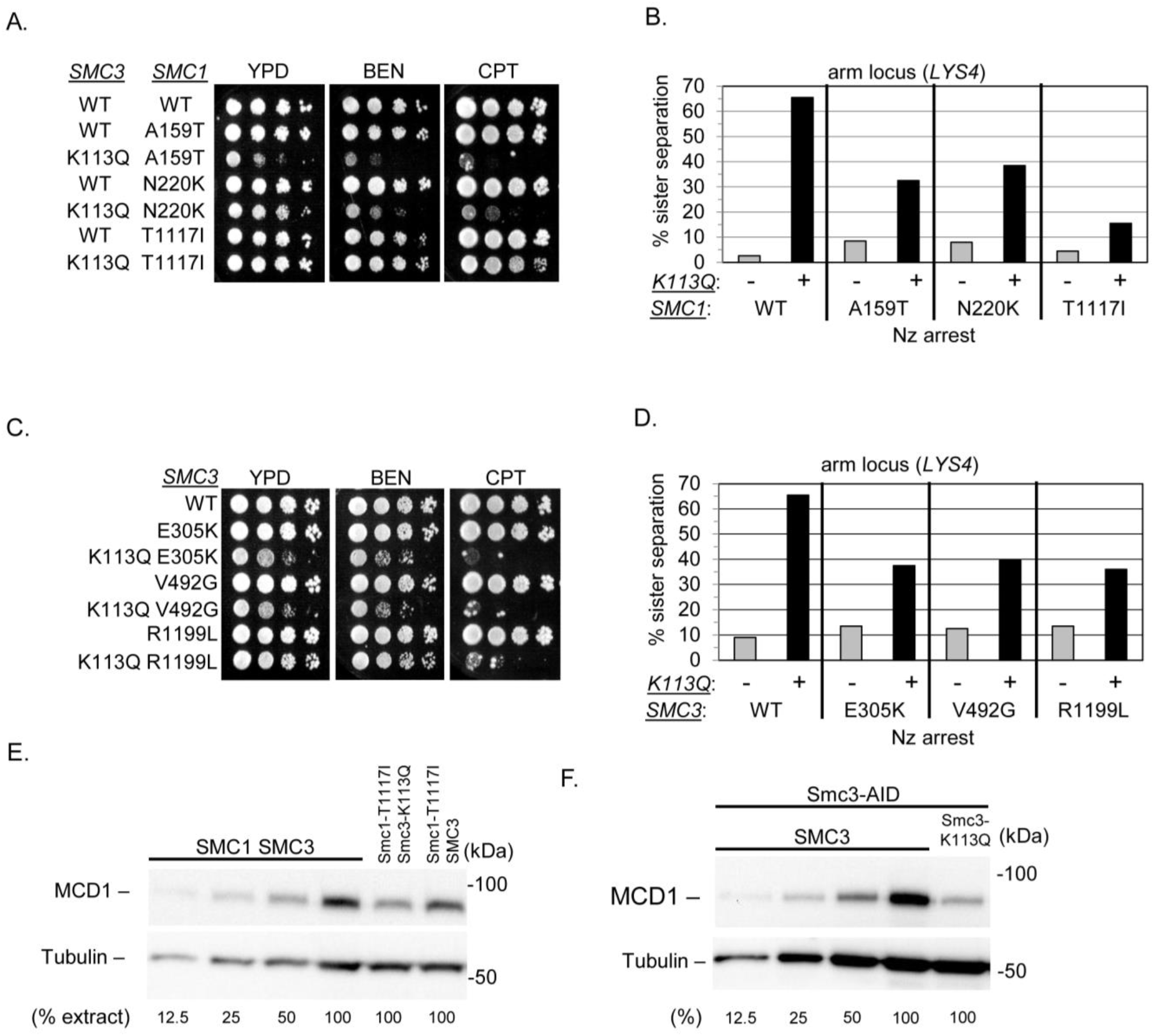
smc1-T1117I is the most robust of the six suppressors of smc3-K113Q inviability. **(A-B)** Comparison of the three SMC1 suppressor mutations shows only smc1-T1117I restores nearly wild-type function to smc3-K113Q cohesin. **(A)** The smc1-T1117I smc3-K113Q double mutant exhibits growth and drug resistance similar to wild-type whereas smc1-A159T smc3-K113Q and smc1-N220K smc3-K113Q exhibit growth defects and drug sensitivity. Haploid wild-type (WT) (VG4012-2C), smc1-A159T (VG4004-7A), smc3-K113Q smc1-A159T (VG4008-2B), smc1-N220K (VG4005-10A), smc3-K113Q smc1-N220K (VG4009-6B), smc1-T1117I (VG4006-13A) and smc3-K113Q smc1-T1117I (VG4010-8B) strains were grown and diluted as described in Figure 1C then plated on YPD alone or containing benomyl [BEN] (10mg/ml) or camptothecin [CPT] (15mg/ml) and incubated for 3d at 23°C, 4d at 23°C or 3d at 30°C respectively. **(B)** smc1-T1117I strongly suppresses cohesion defect of the smc3-K113Q mutant. Haploid strains in (A) were synchronously arrested in mid-M phase and cohesion loss assessed at a chromosome arm site (LYS4) as described in Figure 1D. **(C-D)** Comparison of the three smc3 suppressors of smc3-K113Q shows they all exhibit some defects. **(C)** The three smc3 suppressor smc3-K113Q double mutants exhibit growth defects and drug sensitivity. Haploid wild-type (VG3986-2A), smc3-E305K (VG3997-4C), smc3-K113Q smc3-E305K (VG3998-5B), rsmc3-V492G (VG4000-1A), smc3-K113Q smc3-V492G (VG4001-1B), smc3-R1199L (VG4002-4A), smc3-K113Q smc3-R1199L (VG4003-4B), were grown and plated as described in (A). **(D)** The three smc3 suppressors partially suppress the cohesion defect of the smc3-K113Q mutant. Haploid strains in (C) were synchronously arrested in mid-M phase and cohesion loss assessed at a chromosome arm site (LYS4) as described in Figure 1D. **(E)** Mcd1p levels are restored to approximately 50% of wildtype in the smc1-T1117I smc3-K113Q double mutant. Protein extracts (TCA lysed) from 2 OD aliquots of mid-M phase arrested cells from (C) were analyzed by Western blot. Mcd1p protein levels were monitored using rabbit antibodies against Mcd1p (αMCD1) and rabbit antibodies against tubulin (αTUB2) served as a loading control. **(F)** Mcd1p levels in mid-M phase cells are reduced when acetyl-mimic mutant smc3-K113Q is the sole Smc3p present. Protein extracts (TCA lysed) from 2 OD aliquots of mid-M phase arrested cells from Figure 3B were analyzed by Western blot. Mcd1p protein levels were monitored using rabbit antibodies against Mcd1p (αMCD1) and rabbit antibodies against tubulin (αTUB2) served as a loading control.

**Supplemental Figure 4.**
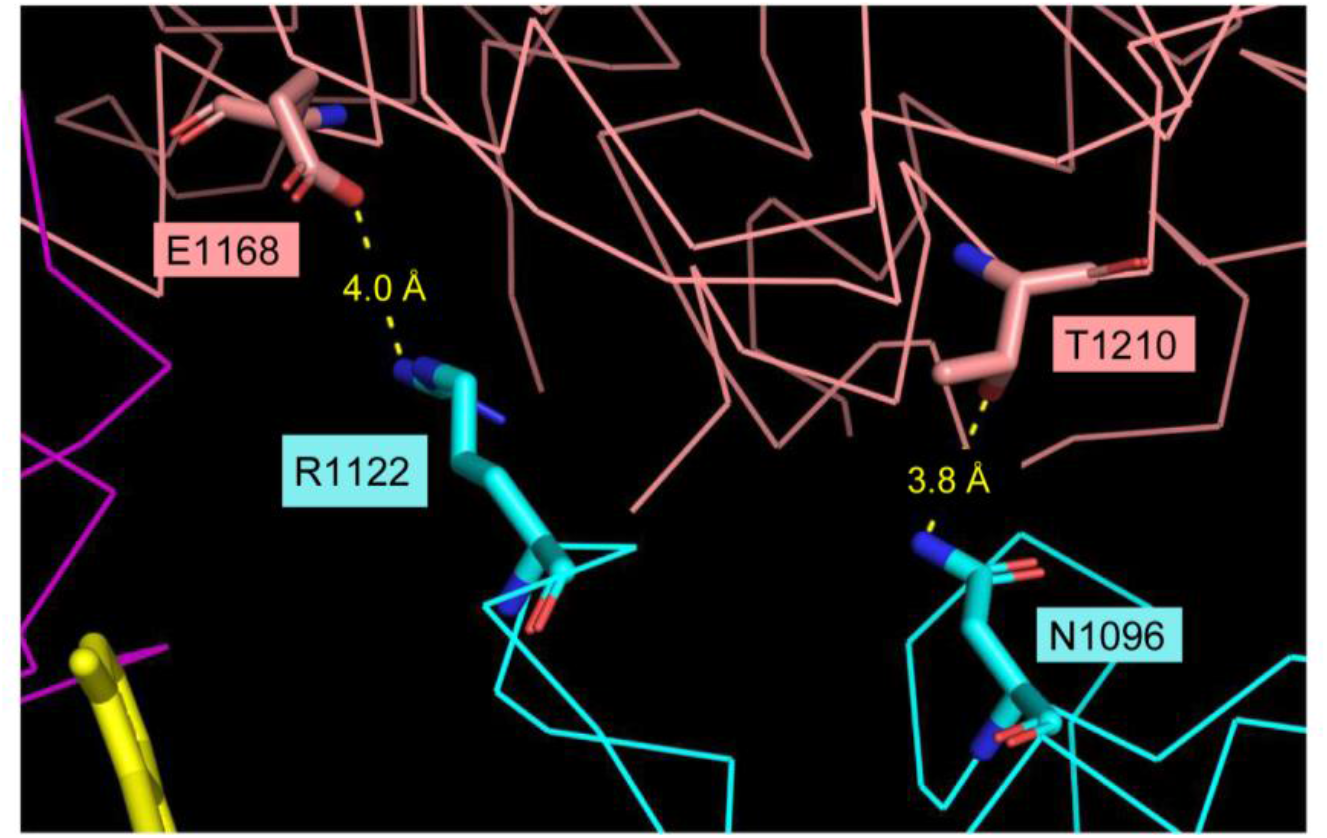
Cryo-EM structure of yeast cohesin and Scc2p with putative salt bridges between Smc1p (cyan) at the Smc1p-T1117 region where it interfaces Scc2p (salmon). Distance measurements indicate spacing between Smc1p-R1122 and Scc2p-E1168 and between Smc1p-N1096 and Scc2p-T1210.

**Supplemental Figure 5.**
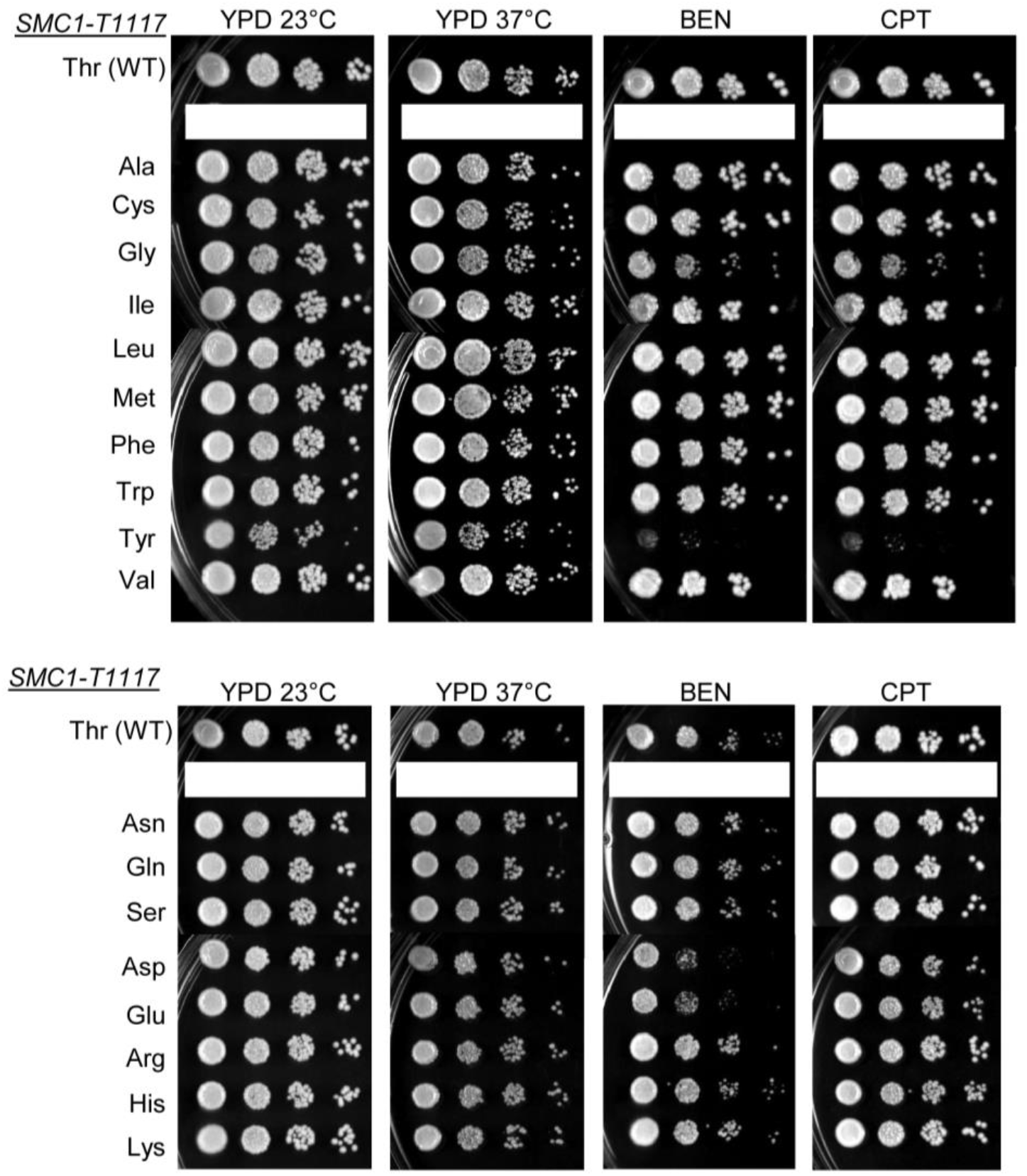
Random mutagenesis of the smc1-T1117 residue in a wild-type background reveals that 19 of the 20 amino acids can support cell viability. CRISPR was used to insert random substitutions of smc1-T1117 residues in a haploid wild-type strain (VG3620-4C) as described in Materials and Methods. Transformant colonies were PCR sequenced to determine which smc1-T1117 substitutions are viable. 19 amino acids were identified, and only proline could not be found. A specific CRISPR to generate Smc1-T1117P confirmed that proline at this residue cannot support viability. Single colonies containing each different amino acid were grown and diluted as described in Figure 1C then plated on YPD and incubated 3d at 23°C or 2d at 37°C or on YPD containing 12.5mg/ml benomyl [BEN] or (25mg/ml camptothecin [CPT] and incubated for 4d at 23°C. All T1117 substitutions grew well at 23°C and 37°C and most were drug resistant. The tyrosine substitution was sensitive to both BEN and CPT whereas aspartic acid and glutamic acid were only BEN sensitive. Plates were electronically rearranged for ease of display.

**Supplemental Figure 6.**
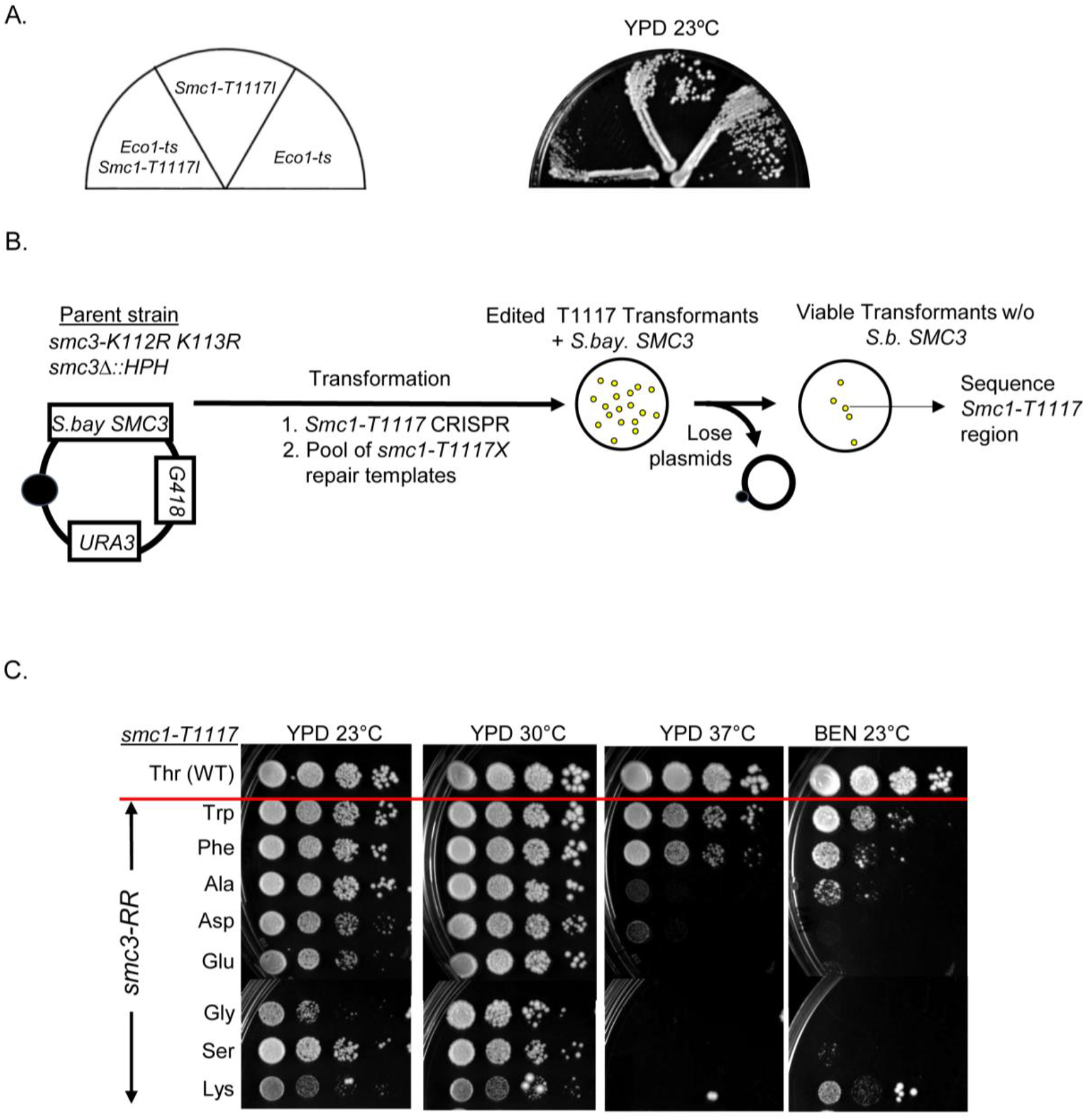
Substitutions at smc1-T1117 can suppress an Smc3 acetyl-null (K112R K113R). **(A)** smc1-T1117I hinders the growth of eco1-ts (ctf7-203) at 23°C. An eco1-203 strain (VG3223-12B), smc1-T1117I eco1-203 strain and an smc1-T1117I (VG4006-13A) strain were dilution streaked on YPD plates and grown at the 23°C for 4 days. **(B)** Schematic of the screen to assess whether any residues at smc1-T1117 can suppress smc3-K112R K113R (RR). Haploid strain (VG4144-5C), containing smc3-K112R, K113R (RR) has the sole S. cerevisiae SMC3 and S. Bayanus SMC3 on a CEN URA3 G418 plasmid (pFC3). CRISPR was used to insert random residues at smc1-T1117. Transformants that are viable bearing only smc3-K113Q as the sole Smc3p in cells were PCR sequenced to identify the smc1-T1117 residue that suppresses smc3-RR inviability. **(C)** smc1-T1117 residues that suppress the lethality of smc3-K112R, K113R (RR) exhibit different growth and drug sensitivity. Haploid wild-type (VG3620-4C) and various smc3-RR strains bearing suppressor mutations at the T1117 residue, T1117W (Trp), T1117F (Phe), T1117A (Ala), T1117E (Asp), T1117E (Glu), T1117G (Gly), T1117S (Ser) and T1117K (Lys), were grown to saturation at 23°C, and then plated at 10-fold serial dilution on YPD and incubated at 23°C 4d, 30°C 3d or 37°C 3d or on YPD containing 10mg/ml benomyl and incubated 23°C 5d (BEN 23°C).

**Supplemental Figure 7.**
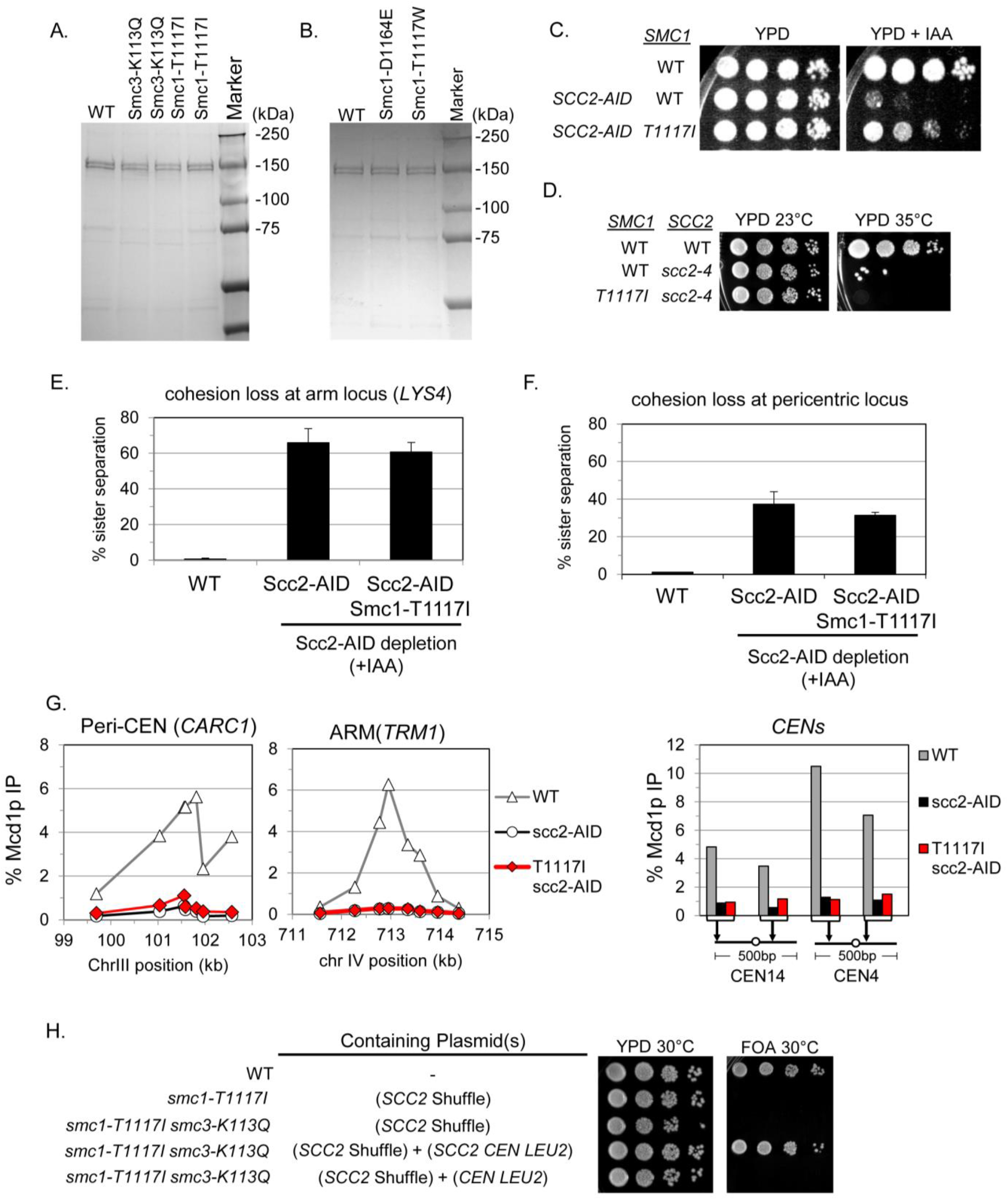
**(A)** Coomassie stain of purified and normalized concentration of wild-type (WT) and mutant cohesin complexes presented in Figure 7A. **(B)** Coomassie stain of purified and normalized concentration of wild-type (WT) and mutant cohesin complexes presented in Figure 7B. **(C)** smc1-T1117I weakly compensates for reduced Scc2p but does not replace Scc2 function. smc1-T1117I suppresses the inviability of scc2-AID depletion. Haploid wild-type (VG3620-4C), scc2-AID (VG3630-7A) and smc1-T1117I scc2-AID (VG4146-1C) strains were grown and diluted as described in Figure 1C then plated on YPD alone or containing IAA then incubated at 23°C for 3d. **(D)** smc1-T1117I cannot suppress a temperature sensitive Scc2p. Haploid wild-type (VG3620-4C), scc2-4 (VG3308-9A) and double mutant smc1-T1117I scc2-4 (VG4162-2A) were grown at 23°C to saturation, plated at 10-fold serial dilutions on YPD and incubated at 23°C or 35°C for 3 days. **(E-F)** smc1-T1117I fails to suppress the cohesion defect of scc2-AID depleted cells. **(E)** Cohesion loss at a chromosomal arm locus (LYS4). Strains in (A) were in synchronously arrested mid-M phase under conditions that deplete Scc2p-AID and cohesion loss at LYS4 as described in Figure 1D. **(F)** Cohesion loss at a CEN-proximal locus. Haploid wild-type (TE631), scc2-AID (VG4163-3A) and smc1-T1117I scc2-AID (VG4164-4B) were synchronously arrested in mid-M phase and cohesion loss at a CEN-proximal locus assessed as described in Figure 1D. **(G)** smc1-T1117I requires Scc2p to load cohesin on chromosomes. Aliquots of mid-M phase arrested cells from (E) were fixed and processed for ChIP, then the level of cohesin bound to chromosomes determined as described in Figure 1E. Left panel is chromosome IV arm CAR region (TRM1), middle panel is chromosome III peri-centric region (CARC1), and right panel are regions immediately adjacent to CEN14 and CEN4. **(H)** smc1-T1117I mutation is unable to bypass the need for SCC2 in either a wild-type background or smc3-K113Q background. Plasmid shuffle assay with wild-type (VG3620-4C), smc1-T1117I scc2Δ + pVG587 (SCC2 shuffle plasmid) (VG4210-1D), smc1-T1117I smc3-K113Q scc2Δ + pVG587 (VG4215-1B), smc1-T1117I smc3-K113Q scc2Δ + pVG587 & p3555 (SCC2 CEN LEU2) (VG4215-1B + p3555), and smc1-T1117I smc3-K113Q scc2Δ + pVG587 & pRS315 (CEN LEU2) (VG4215-1B + pRS315). Strains were grown and diluted as in Figure 1C then plated on YPD and 5-FOA media and incubated for 2 days at 30°C.

