## Supplemental Table 1 - Strain List for "A model for Scc2p Stimulation of Cohesin’s ATPase and its Inhibition by Acetylation of Smc3p"

Supplemental Table 1 – Strain Table

| Strain | Genotype | Background |
|---|---|---|
| VG3919-3C | MATa SMC3-LEU2:leu2-3,112 smc3-N607-3V5-AID TIR1-CaTRP1 LacO-NAT::lys4 GFPLacI-HIS3:his3-11,15 ura3-52 bar1 GAL+ | A346A |
| VG3981-6B | MATa SMC3-K113Q-LEU2:leu2-3,112 smc3-N607-3V5-AID TIR1-CaTRP1<br>LacO-NAT::lys4 GFPLacI-HIS3:his3-11,15 ura3-52 bar1 GAL+ | A346A |
| VG4052-3A | MATa pGAL-SCC2/4-URA3:ura3-52 TIR1-CaTRP1 smc3-K113Q-LEU2:leu2-3,112 smc3-N607-3V5-AID LacO-NAT::lys4 GFPLacI-HIS3:his3-11,15 bar1 GAL+ | A346A |
| VG3651-3D | MATa SMC3-N607-3V5-AID TIR1-CgTRP1 LacO-NAT::lys4 leu2-3,112 ura3-52 GFPLacI-HIS3:his3-11,15 bar1 GAL+ | A346A |
| JL17A | MATa SMC3-LEU2:leu2-3,112 smc3-N607-3V5-AID TIR1-CaTRP1 LacO:10kb-CEN4-NAT GFPLacI-HIS3:his3-11,15 ura3-52 bar1 GAL+ | A346A |
| JL16A | MATa SMC3-N607-3V5-AID TIR1-CgTRP1 LacO:10kb-CEN4-NAT leu2-3,112 ura3-52 GFPLacI-HIS3:his3-11,15 bar1 GAL+ | A346A |
| JL14A | MATa SMC3-K113Q-LEU2:leu2-3,112 smc3-N607-3V5-AID TIR1-CaTRP1 LacO:10kb-CEN4-NAT GFPLacI-HIS3:his3-11,15 URA3:ura3-52 bar1 GAL+ | A346A |
| JL15A | MATa pGAL-SCC2/4-URA3:ura3-52 TIR1-CaTRP1 smc3-K113Q-LEU2:leu2-3,112 smc3-N607-3V5-AID LacO:10kb-CEN4-NAT GFPLacI-HIS3:his3-11,15 bar1 GAL+ | A346A |
| KB61A | MATa TIR1-CgTRP1 LacO-NAT::lys4 GFPLacI-HIS3:his3-11,15 URA3:ura3-52 leu2-3,112 bar1 GAL+ | A346A |
| VG3633-2D | MATa ECO1-3V5-AID2-G418 TIR1-CaTRP1 LacO-NAT::lys4 leu2-3,112 GAL+ pHIS3-GFPLacI-HIS3:his3-11,15 bar1 ura3-52 | A346A |
| KB50A | MATa ECO1-3V5-AID2-G418 TIR1-CaTRP1 LacO-NAT::lys4 pGAL-SCC2,SCC4-LEU2:leu2-3,112 GAL+ pHIS3-GFPLacI-HIS3:his3-11,15 bar1 ura3-52 | A346A |
| VG3862-1A | MATa PDS5-3V5-AID2F:G418 TIR1-CaTRP1 LacO-NAT::lys4 GFPLacI-HIS3:his3-11,15 leu2-3,112 ura3-52 bar1 GAL+ | A346A |
| KB46A | MATa PDS5-3V5-AID2F:G418 TIR1-CaTRP1 LacO-NAT::lys4 GFPLacI-HIS3:his3-11,15 leu2-3,112 pGAL-SCC2,SCC4-URA3:ura3-52 bar1 GAL+ | A346A |
| KB47A | MATa SMC3-N607-3V5-AID TIR1-CgTRP1 LacO-NAT::lys4 leu2-3,112 URA3-pGAL1,10-SCC2, SCC4::ura3-52 GFPLacI-HIS3:his3-11,15 bar1 GAL+ | A346A |

| | | |
|---|---|---|
| VG3969-14C | MATa smc3Δ::HPH smc3-K113Q-LEU2:leu2-3,112 LacO-NAT::lys4 GFPLacI-TRP1:his3-11,15 trp1-1 ura3-52 bar1 GAL+ w/pFC3 | A346A |
| VG4012-2C | MATa SMC1-LEU2:leu2-3,112 smc1Δ::HPH LacO-NAT::lys4 GFPLacI-HIS3:his3-11,15 trp1-1 ura3-52 bar1 GAL+ | A346A |
| VG4005-10A | MATa smc1-N220K-LEU2:leu2-3,112 smc1Δ::HPH LacO-NAT::lys4 GFPLacI-HIS3:his3-11,15 trp1-1 ura3-52 bar1 GAL+ | A346A |
| VG4009-6B | MATa smc3-K113Q smc1-N220K-LEU2:leu2-3,112 smc1Δ::HPH LacO-NAT::lys4 GFPLacI-HIS3:his3-11,15 trp1-1 ura3-52 bar1 GAL+ | A346A |
| VG4006-13A | MATa smc1-T1117I-LEU2:leu2-3,112 smc1Δ::HPH LacO-NAT::lys4 GFPLacI-HIS3:his3-11,15 trp1-1 ura3-52 bar1 GAL+ | A346A |
| VG4010-8B | MATa smc3-K113Q smc1-T1117I-LEU2:leu2-3,112 smc1Δ::HPH LacO-NAT::lys4 GFPLacI-HIS3:his3-11,15 trp1-1 ura3-52 bar1 GAL+ | A346A |
| 3620-4C | MATa TIR1-CgTRP1 LacO-NAT::lys4 GFPLacI-HIS3:his3-11,15 leu2-3,112 ura3-52 bar1 GAL+ | A346A |
| VG4004-7A | MATa smc1-A159T-LEU2:leu2-3,112 smc1Δ::HPH LacO-NAT::lys4 GFPLacI-HIS3:his3-11,15 trp1-1 ura3-52 bar1 GAL+ | A346A |
| VG4008-2B | MATa smc3-K113Q smc1-A159T-LEU2:leu2-3,112 smc1Δ::HPH LacO-NAT::lys4 GFPLacI-HIS3:his3-11,15 trp1-1 ura3-52 bar1 GAL+ | A346A |
| VG3986-2A | MATa SMC3-LEU2:leu2-3,112 smc3Δ::HPH LacO-NAT::lys4 GFPLacI-TRP1:his3-11,15 trp1-1 ura3-52 bar1 GAL+ | A346A |
| VG3997-4C | MATa smc3Δ::HPH smc3-E305K-LEU2:leu2-3,112 LacO-NAT::lys4 GFPLacI-TRP1:his3-11,15 trp1-1 ura3-52 bar1 GAL+ | A346A |
| VG3998-5B | MATa smc3Δ::HPH smc3-K113Q,E305K-LEU2:leu2-3,112 LacO-NAT::lys4 GFPLacI-TRP1:his3-11,15 trp1-1 ura3-52 bar1 GAL+ | A346A |
| VG4000-1A | MATa smc3-V492G-LEU2:leu2-3,112 smc3Δ::HPH LacO-NAT::lys4 GFPLacI-TRP1:his3-11,15 trp1-1 ura3-52 bar1 GAL+ | A346A |
| VG4001-1B | MATa smc3-K113Q,V492G-LEU2:leu2-3,112 smc3Δ::HPH LacO-NAT::lys4 GFPLacI-TRP1:his3-11,15 trp1-1 ura3-52 bar1 GAL+ | A346A |
| VG4002-4A | MATa smc3-R1199L-LEU2:leu2-3,112 smc3Δ::HPH LacO-NAT::lys4 GFPLacI-TRP1:his3-11,15 trp1-1 ura3-52 bar1 GAL+ | A346A |
| VG4003-4B | MATa smc3-K113Q,R1199L-LEU2:leu2-3,112 smc3Δ::HPH LacO-NAT::lys4 GFPLacI-TRP1:his3-11,15 trp1-1 ura3-52 bar1 GAL+ | A346A |
| VG3223-12B | MATa ctf7-203 LacO-NAT::lys4 pHIS3-GFPLacI-HIS3:his3-11,15 GAL+ trp1-1 leu2-3,112 ura3-52 bar1 | A346A |
| KB88 | MATa ctf7-203 smc1-T1117I LacO-NAT::lys4 pHIS3-GFPLacI-HIS3:his3-11,15 GAL+ trp1-1 leu2-3,112 ura3-52 bar1 | A346A |

| | | |
|---|---|---|
| VG4147-14C | MATa smc3-K113Q smc1-T1117V-LEU2:leu2-3,112 smc1Δ::HPH LacO-NAT::lys4 GFPLacI-HIS3:his3-11,15 trp1-1 ura3-52 bar1 GAL+ | A346A |
| VG3633-3D | MATa ECO1-3V5-AID2-G418 TIR1-CaTRP1 LacO-NAT::lys4 leu2-3,112 GAL+ pHIS3-GFPLacI-HIS3:his3-11,15 bar1 ura3-52 | A346A |
| KB118E | MATa smc1-T1117I ECO1-3V5-AID2-G418 TIR1-CaTRP1 LacO-NAT::lys4 leu2-3,112 GAL+pHIS3-GFPLacI-HIS3:his3-11,15 bar1 ura3-52 | A346A |
| JL11A | MATa smc1-T1117I TIR1-CgTRP1 LacO-NAT::lys4 GFPLacI-HIS3:his3-11,15 leu2-3,112 ura3-52 bar1 GAL+ | A346A |
| VG3620-4C Ala | MATa smc1-T1117A TIR1-CgTRP1 LacO-NAT::lys4 GFPLacI-HIS3:his3-11,15 leu2-3,112 ura3-52 bar1 GAL+ | A346A |
| VG3620-4C Cys | MATa smc1-T1117C TIR1-CgTRP1 LacO-NAT::lys4 GFPLacI-HIS3:his3-11,15 leu2-3,112 ura3-52 bar1 GAL+ | A346A |
| VG3620-4C Gly | MATa smc1-T1117G TIR1-CgTRP1 LacO-NAT::lys4 GFPLacI-HIS3:his3-11,15 leu2-3,112 ura3-52 bar1 GAL+ | A346A |
| VG3620-4C Ile | MATa smc1-T1117I TIR1-CgTRP1 LacO-NAT::lys4 GFPLacI-HIS3:his3-11,15 leu2-3,112 ura3-52 bar1 GAL+ | A346A |
| VG3620-4C Leu | MATa smc1-T1117L TIR1-CgTRP1 LacO-NAT::lys4 GFPLacI-HIS3:his3-11,15 leu2-3,112 ura3-52 bar1 GAL+ | A346A |
| VG3620-4C Met | MATa smc1-T1117M TIR1-CgTRP1 LacO-NAT::lys4 GFPLacI-HIS3:his3-11,15 leu2-3,112 ura3-52 bar1 GAL+ | A346A |
| VG3620-4C Phe | MATa smc1-T1117F TIR1-CgTRP1 LacO-NAT::lys4 GFPLacI-HIS3:his3-11,15 leu2-3,112 ura3-52 bar1 GAL+ | A346A |
| VG3620-4C Trp | MATa smc1-T1117W TIR1-CgTRP1 LacO-NAT::lys4 GFPLacI-HIS3:his3-11,15 leu2-3,112 ura3-52 bar1 GAL+ | A346A |
| VG3620-4C Tyr | MATa smc1-T1117Y TIR1-CgTRP1 LacO-NAT::lys4 GFPLacI-HIS3:his3-11,15 leu2-3,112 ura3-52 bar1 GAL+ | A346A |
| VG3620-4C Val | MATa smc1-T1117V TIR1-CgTRP1 LacO-NAT::lys4 GFPLacI-HIS3:his3-11,15 leu2-3,112 ura3-52 bar1 GAL+ | A346A |
| VG3620-4C Asn | MATa smc1-T1117N TIR1-CgTRP1 LacO-NAT::lys4 GFPLacI-HIS3:his3-11,15 leu2-3,112 ura3-52 bar1 GAL+ | A346A |
| VG3620-4C Gln | MATa smc1-T1117Q TIR1-CgTRP1 LacO-NAT::lys4 GFPLacI-HIS3:his3-11,15 leu2-3,112 ura3-52 bar1 GAL+ | A346A |
| VG3620-4C Ser | MATa smc1-T1117S TIR1-CgTRP1 LacO-NAT::lys4 GFPLacI-HIS3:his3-11,15 leu2-3,112 ura3-52 bar1 GAL+ | A346A |
| VG3620-4C Asp | MATa smc1-T1117D TIR1-CgTRP1 LacO-NAT::lys4 GFPLacI-HIS3:his3-11,15 leu2-3,112 ura3-52 bar1 GAL+ | A346A |

| | | |
|---|---|---|
| VG3620-4C Glu | MATa smc1-T1117E TIR1-CgTRP1 LacO-NAT::lys4 GFPLacI-HIS3:his3-11,15 leu2-3,112 ura3-52 bar1 GAL+ | A346A |
| VG3620-4C Arg | MATa smc1-T1117R TIR1-CgTRP1 LacO-NAT::lys4 GFPLacI-HIS3:his3-11,15 leu2-3,112 ura3-52 bar1 GAL+ | A346A |
| VG3620-4C His | MATa smc1-T1117H TIR1-CgTRP1 LacO-NAT::lys4 GFPLacI-HIS3:his3-11,15 leu2-3,112 ura3-52 bar1 GAL+ | A346A |
| VG3620-4C Lys | MATa smc1-T1117K TIR1-CgTRP1 LacO-NAT::lys4 GFPLacI-HIS3:his3-11,15 leu2-3,112 ura3-52 bar1 GAL+ | A346A |
| VG3630-7A | MATa SCC2-3V5-AID2-G418 TIR1-CgTRP1 LacO-NAT::lys4 leu2-3,112 GAL+ GFPLacI-HIS3:his3-11,15 bar1 ura3-52 | A346A |
| VG4146-1C | MATa smc1-T1117I SCC2-3V5-AID2-G418 TIR1-CgTRP1 LacO-NAT::lys4 GFPLacI-HIS3:his3-11,15 leu2-3,112 ura3-52 bar1 GAL+ | A346A |
| TE631 | MATa LacO-NAT:10KB-CEN4 TIR1-CaTRP1 leu2-3,112 GFPLacI-HIS3:his3-11,15 ura3-52 bar1 GAL+ | A346A |
| VG4163-3A | MATa SCC2-3V5-AID2-G418 TIR1-CgTRP1 LacO-NAT:10kb-CEN4 leu2-3,112 GFPLacI-HIS3:his3-11,15 ura3-52 bar1 GAL+ | A346A |
| VG4164-4B | MATa smc1-T1117I SCC2-3V5-AID2-G418 TIR1-CgTRP1 LacO-NAT:10kb-CEN4 GFPLacI-HIS3:his3-11,15 leu2-3,112 ura3-52 bar1 GAL+ | A346A |
| VG3308-9A | MATa scc2-4 trp1-1 leu2-3,112 ura3-52 GFPLacI-HIS3:his3-11,15 bar1 GAL+ | A346A |
| VG4162-2A | MATa smc1-T1117I scc2-4 LacO-NAT::lys4 GFPLacI-HIS3:his3-11,15 trp1-1 leu2-3,112 ura3-52 bar1 GAL+ | A346A |
| VG4144-5C | MATa smc3Δ::HPH smc3-K112R,K113R-LEU2:leu2-3,112 GFPLacI-TRP1:his3-11,15 trp1-1 LacO-NAT::lys4 ura3-52 bar1 GAL+ w/pFC3 | A346A |
| VG4144-5C Trp | MATa smc1-T1117W smc3Δ::HPH smc3-K112R,K113R-LEU2:leu2-3,112 GFPLacI-TRP1:his3-11,15 trp1-1 LacO-NAT::lys4 ura3-52 bar1 GAL+ | A346A |
| VG4144-5C Phe | MATa smc1-T1117F smc3Δ::HPH smc3-K112R,K113R-LEU2:leu2-3,112 GFPLacI-TRP1:his3-11,15 trp1-1 LacO-NAT::lys4 ura3-52 bar1 GAL+ | A346A |
| VG4144-5C Ala | MATa smc1-T1117A smc3Δ::HPH smc3-K112R,K113R-LEU2:leu2-3,112 GFPLacI-TRP1:his3-11,15 trp1-1 LacO-NAT::lys4 ura3-52 bar1 GAL+ | A346A |
| VG4144-5C Asp | MATa smc1-T1117D smc3Δ::HPH smc3-K112R,K113R-LEU2:leu2-3,112 GFPLacI-TRP1:his3-11,15 trp1-1 LacO-NAT::lys4 ura3-52 bar1 GAL+ | A346A |

| | | |
|---|---|---|
| VG4144-5C Glu | MATa smc1-T1117E smc3Δ::HPH smc3-K112R,K113R-LEU2:leu2-3,112 GFPLacI-TRP1:his3-11,15 trp1-1 LacO-NAT::lys4 ura3-52 bar1 GAL+ | A346A |
| VG4144-5C Gly | MATa smc1-T1117G smc3Δ::HPH smc3-K112R,K113R-LEU2:leu2-3,112 GFPLacI-TRP1:his3-11,15 trp1-1 LacO-NAT::lys4 ura3-52 bar1 GAL+ | A346A |
| VG4144-5C Ser | MATa smc1-T1117S smc3Δ::HPH smc3-K112R,K113R-LEU2:leu2-3,112 GFPLacI-TRP1:his3-11,15 trp1-1 LacO-NAT::lys4 ura3-52 bar1 GAL+ | A346A |
| VG4144-5C Lys | MATa smc1-T1117K smc3Δ::HPH smc3-K112R,K113R-LEU2:leu2-3,112 GFPLacI-TRP1:his3-11,15 trp1-1 LacO-NAT::lys4 ura3-52 bar1 GAL+ | A346A |
| VG4154-3A | MATa wpl1Δ::HPH smc3-K112R,K113R LacO-NAT::lys4 GFPLacI-HIS3:his3-11,15 trp1-1 leu2-3,112 ura3-52 bar1 GAL+ | A346A |
| VG4153-5C | MATa smc1-D1164E smc3-K112R,K113R TIR1-CgTRP1 LacO-NAT::lys4 GFPLacI-HIS3:his3-11,15 leu2-3,112 ura3-52 bar1 GAL+ | A346A |
| VG4158-9D | MATa smc1-T1117W smc3-K112R,K113R-LEU2:leu2-3,112 smc3Δ::HPH LacO-NAT::lys4 GFPLacI-TRP1:his3-11,15 trp1-1 ura3-52 bar1 GAL+ | A346A |
| VG3991-1A | MATa smc3-K112R,K113R-LEU2:leu2-3,112 SMC3-N607-3V5-AID TIR1-CgTRP1 LacO-NAT::lys4 GFPLacI-HIS3:his3-11,15 ura3-52 bar1 GAL+ | A346A |
| VG4168-7B | MATa smc1-T1117W TIR1-CgTRP1 LacO-NAT::lys4 GFPLacI-HIS3:his3-11,15 leu2-3,112 ura3-52 bar1 GAL+ | A346A |
| VG4138-1A | MATa smc1-D1164E TIR1-CgTRP1 LacO-NAT::lys4 GFPLacI-HIS3:his3-11,15 leu2-3,112 ura3-52 bar1 GAL+ | A346A |
| VG3961-4B | MATa smc3Δ::HPH LacO-NAT::lys4 GFPLacI-TRP1:his3-11,15 trp1-1 leu2-3,112 ura3-52 bar1 GAL+ w/pFC3 | A346A |
| VG3965-1A | MATa smc1Δ::HPH LacO-NAT::lys4 GFPLacI-HIS3:his3-11,15 trp1-1 leu2-3,112 ura3-52 bar1 GAL+ w/pFC1 | A346A |
| KB58A | MATa pGAL1,10-GAL4, SMC1-PK3-ADE2:ade2-1, pGAL1,10-SCC3-MYC-URA3:ura3, pGAL1,10 SMC3,MCD1-3C-3xStrepII-TRP1:trp1, can1-100, leu2-3,112, his3, GAL, psi+, pep4Δ::HIS3 wpl1Δ::LEU eco1Δ::KANMX6 | K699 |
| KB140A | MATa pGAL1,10-GAL4, SMC1-PK3-ADE2:ade2-1, pGAL1,10-SCC3-MYC-URA3:ura3, pGAL1,10 smc3-K113Q, MCD1-3C-3xStrepII-TRP1:trp1, can1-100, leu2-3,112, his3, GAL, psi+, pep4Δ::HIS3 wpl1Δ::LEU eco1Δ::KANMX6 | K699 |
| SX313 | MATa pGAL1,10-GAL4, SMC1-T1117I-PK3-ADE2:ade2-1, pGAL1,10-SCC3-MYC-URA3:ura3, pGAL1,10 smc3-K113Q, MCD1- | K699 |

| | | |
|---|---|---|
| | 3C-3xStrepII-TRP1:trp1, can1-100, leu2-3,112, his3, GAL, psi+, pep4Δ::HIS3 wpl1Δ::LEU eco1Δ::KANMX6 | |
| SX312 | MATa pGAL1,10-GAL4, SMC1-T1117I-PK3-ADE2:ade2-1, pGAL1,10-SCC3-MYC-URA3:ura3, pGAL1,10 SMC3,MCD1-3C-3xStrepII-TRP1:trp1, can1-100, leu2-3,112, his3, GAL, psi+, pep4Δ::HIS3 wpl1Δ::LEU eco1Δ::KANMX6 | K699 |
| SX318 | MATa pGAL1,10-GAL4, SMC1-T1117W-PK3-ADE2:ade2-1, pGAL1,10-SCC3-MYC-URA3:ura3, pGAL1,10 SMC3,MCD1-3C-3xStrepII-TRP1:trp1, can1-100, leu2-3,112, his3, GAL, psi+, pep4Δ::HIS3 wpl1Δ::LEU eco1Δ::KANMX6 | K699 |
| SX316 | MATa pGAL1,10-GAL4, SMC1-D1164E-PK3-ADE2:ade2-1, pGAL1,10-SCC3-MYC-URA3:ura3, pGAL1,10 SMC3,MCD1-3C-3xStrepII-TRP1:trp1, can1-100, leu2-3,112, his3, GAL, psi+, pep4Δ::HIS3 wpl1Δ::LEU eco1Δ::KANMX6 | K699 |
| SX305 | MATa lys2 pep4::HIS3 ade2-1::ADE2-pGAL-GAL4 trp1Δ2 leu2-3, 112 ura3-52 ade2-1 can1-100 bar1::hisG w/[ 2u URA3 pGAL1,10-(Scc2-MYC-Strept)-(Scc4) ] | W303 |
| VG4210-1D | MATa smc1-T1117I scc2Δ::G418 TIR1-CgTRP1 LacO-NAT::lys4 GFPLacI-HIS3:his3-11,15 leu2-3,112 ura3-52 bar1 GAL+ w/pVG587 (SCC2 URA3 CEN) | A346A |
| VG4215-1B | MATa smc1-T1117I smc3-K113Q scc2Δ::G418 TIR1-CgTRP1 LacO-NAT::lys4 GFPLacI-HIS3:his3-11,15 leu2-3,112 ura3-52 bar1 GAL+ w/pVG587 (SCC2 URA3 CEN) | A346A |
| VG4215-1B + p3555 | MATa smc1-T1117I smc3-K113Q scc2Δ::G418 TIR1-CgTRP1 LacO-NAT::lys4 GFPLacI-HIS3:his3-11,15 leu2-3,112 ura3-52 bar1 GAL+ w/pVG587 (SCC2 URA3 CEN) & p3555 (SCC2 LEU2 CEN) | A346A |
| VG4215-1B + pRS315 | MATa smc1-T1117I smc3-K113Q scc2Δ::G418 TIR1-CgTRP1 LacO-NAT::lys4 GFPLacI-HIS3:his3-11,15 leu2-3,112 ura3-52 bar1 GAL+ w/pVG587 (SCC2 URA3 CEN) & pRS315 (LEU2 CEN) | A346A |
