## Supplemental Table 2 - Plasmid List for "A model for Scc2p Stimulation of Cohesin’s ATPase and its Inhibition by Acetylation of Smc3p"

Supplemental Table 2 – Plasmid Table

### Integrative Plasmids

| Plasmid name | Yeast Genes |
|---|---|
| pVG556 | pGAL-SCC2, pGAL-SCC4 LEU2 |
| pFC5 | pGAL-SCC2 pGAL-SCC4 URA3 |
| pVG444 | SMC1 LEU2 |
| pVG444 A159T | smc1-A159T LEU2 |
| pVG444 N220K | smc1-N220K LEU2 |
| pVG444 T1117I | smc1-T1117I LEU2 |
| pVG419 | SMC3 LEU2 |
| pVG419 K113Q | smc3-K113Q LEU2 |
| pVG419 E305K | smc3-E305K LEU2 |
| pVG419 K113Q, E305K | smc3-K113Q, E305K LEU2 |
| pVG419 V492G | smc3-V492G LEU2 |
| pVG419 K113Q, V492G | smc3-K113Q, V492G LEU2 |
| pVG419 R1199L | smc3-R1199L LEU2 |
| pVG419 K113Q, R1199L | smc3-K113Q, R1199L LEU2 |

### Centromere Plasmids

| Plasmid name | Yeast genes |
|---|---|
| pFC1 | S.bay SMC1 URA3 G418 |
| pFC3 | S.bay SMC3 URA3 G418 |

### CRISPR Plasmids

| Plasmid | Guide Target | Primers | Parent plasmid |
|---|---|---|---|
| pFC8 | Smc1-T1117 | VG774/VG775 | pJR3429 |
| pVG524 | Smc1-K113 | VG727/VG728 | pVG522 |
| pFC9 | Smc1-D1164 | VG766/VG767 | pJR3429 |

### Purification Plasmids

| Purification Plasmids | Yeast Genes |
|---|---|
| pSX103 | pGAL-SMC1-PK3, pGAL-GAL4 ADE2 |
| pSX104 | pGAL-SCC3-MYC URA3 |
| pKB11 | pGAL-SMC3, pGAL-MCD1-3C-3xStrepII TRP1 |
| pKB57 | pGAL-Smc1T1117I-PK3, pGAL-GAL4 ADE2 |
| pKB58 | pGAL-Smc1-T1117W-PK3, pGAL-GAL4 ADE2 |
| pKB47 | pGAL-Smc3-K113Q, pGAL-MCD1-3C-3xStrepII TRP1 |
| pKB49B | pGAL-Smc3-K38I, pGAL-MCD1-3C-3xStrepII TRP1 |

### 2-micron Plasmid

| Purification Plasmids | Yeast Genes |
|---|---|
| GC1206 | pGAL-SCC4, pGAL-SCC2-MYC3-3xStrepII URA3 |
