## Supplemental Table 3 - Primer List for "A model for Scc2p Stimulation of Cohesin’s ATPase and its Inhibition by Acetylation of Smc3p"

| **CRISPR Guide Primers**<br>(upper case bases are genomic sequences; lower case are bases to anneal to BsmBI ends) | | |
|---|---|---|
| Primer | Sequence (5'-->3') | Plasmid Name |
| VG774 | gactttGAATCTTTTAAGAGGCGGAG | pFC8 |
| VG775 | aaacCTCCGCCTCTTAAAAGATTCaa | pFC8 |
| VG727 | gactttAAGAACAGTAGGGCTGAAGA | pVG524 |
| VG728 | aaacTCTTCAGCCCTACTGTTCTTaa | pVG524 |
| VG766 | gactttCTGGACGTTAGTAATGTCTA | pFC9 |
| VG767 | aaacTAGACATTACTAACGTCCAGaa | pFC9 |

| **CRISPR Repair Template Primers** | | |
|---|---|---|
| Primer | Sequence (changed codon for mutation is shown in red) | Mutation Introduced |
| VG749 | CTT TCG AGA GGA GAT GAC GAA GTG ACC ATT AGA AGA ACA GTA GGG CTG AAG cAG GAT GAC | Smc3-K113Q (sense strand) |
| VG750 | CTA TGT CCC CTT TGG TCA CGT TTC TGT CAT TTA ATT GAT AGT CAT CCT gCT TCA GCC CTA | Smc3-K113Q (anti-sense strand) |
| VG868 | GAA GAT GAA CCG TTC AAT GCG GGA ATC AAA TAT CAT GCC AtT CCt CCT CTT AAA AGA TTC | Smc1-T1117I (sense strand) |
| VG869 | TTC ACC ACC AGA AAG ATA TTC CAT GTC TTT GAA TCT TTT AAG AGG aGG AaT GGC ATG ATA | Smc1-T1117I (anti-sense strand) |
| VG870 | AC CAG CCT AGT CCC TTC TTC GTG CTG GAC GAA GTG GAC GCA GCC CTA GAa ATT ACT AAC G | Smc1-D1164E (sense strand) |
| VG871 | GG ATT ACG GTG CCT TCT TAT ATA GGC AGC AAT TCT CTG GAC GTT AGT AAT tTC TAG GGC T | Smc1-D1164E (anti-sense strand) |
| VG1002 | GAA GAT GAA CCG TTC AAT GCG GGA ATC AAA TAT CAT GCC NNK CCt CCT CTT AAA AGA TTC | Smc1-T1117X (sense strand) |
| VG1003 | TTC ACC ACC AGA AAG ATA TTC CAT GTC TTT GAA TCT TTT AAG AGG aGG KNN GGC ATG ATA | Smc1-T1117X (anti-sense strand) |
| VG1016 | GAA GAT GAA CCG TTC AAT GCG GGA ATC AAA TAT CAT GCC tgg CCt CCT CTT AAA AGA TTC | Smc1-T1117W (sense strand) |
| VG1017 | TTC ACC ACC AGA AAG ATA TTC CAT GTC TTT GAA TCT TTT AAG AGG aGG cca GGC ATG ATA | Smc1-T1117W (anti-sense strand) |

| qPCR Primers | | |
|---|---|---|
| BR463 | CATGATTCGCCGGGTAAATA | CEN4 left arm (4A F) |
| BR464 | GCACTAGCCAATTTAGCACTTC | CEN4 left arm (4A R) |
| BR465 | AAAATGCCGAGGCTTTCATA | CEN4 right arm (4B F) |
| BR466 | TGACGATAAAACCGGAAGGA | CEN4 right arm (4B R) |
| TE442 | TTAAAGCGGCTGAGTATGGC | CEN14 left arm (14A F) |
| TE443 | TTTCCTCCATTGCTCTCTACGG | CEN14 left arm (14A R) |
| TE446 | ACTAAAAGTGCCCCAAACGG | CEN14 right arm (14B F) |
| TE447 | AGGAGCAGGGTAGCATAAACC | CEN14 right arm (14B R) |
| VG641 | AAA GAA GCA GGG GTA GAG AAG C | TRM1 (CEN distal; T0 F) |
| VG642 | ATC AGC AGC GGT GAT TAC AC | TRM1 (CEN distal; T0 R) |
| VG625 | CCA GCC AGA TAT TAT GGG CAA G | TRM1 (CEN distal; T1 F) |
| VG626 | TCC TAG ACC TGG TGG AAA AAG C | TRM1 (CEN distal; T1 R) |
| VG627 | CTT ATA GTT CCC AAG GCA TCC C | TRM1 (CEN distal; T2 F) |
| VG628 | CCA AAC TCG TTG TTC TCG ATC C | TRM1 (CEN distal; T2 R) |
| VG629 | TCT TCG TGC GCG AGG ATA TG | TRM1 (CEN distal; T3 F) |
| VG630 | CGA ACA TTT CCG GAC AAT TGC | TRM1 (CEN distal; T3 R) |
| VG633 | CCA ATC GTA TAA CGG AGC ATT GG | TRM1 (CEN distal; T5 F) |
| VG634 | TGG TGC CAG AAG ATA TCA ACG | TRM1 (CEN distal; T5 R) |
| VG637 | GCG CGA TAC CAT TCA GAA CAT C | TRM1 (CEN distal; T7 F) |
| VG638 | TTA AAG TGG GCC CAA GAC CAG | TRM1 (CEN distal; T7 R) |
| VG639 | GGG CAT CAC CTT TTC GTA AGC | TRM1 (CEN distal; T8 R) |
| VG640 | TGA TCC ACC TGT CAT TTC GC | TRM1 (CEN distal; T8 R) |
| VG643 | ACC CTT CTG TTC CAG TTT GC | TRM1 (CEN distal; T9 F) |

| | | |
|---|---|---|
| VG644 | GTT GCC TCC GGA GCA AAT TC | TRM1 (CEN distal; T9 R) |
| TE367 | AAA GGT GCC CCA AGA AAA GG | CARC1 (pericentric; 1 F) |
| TE368 | AGC ACT TTA CTC GCT TGT GG | CARC1 (pericentric; 1 R) |
| TE310 | TAA AGC ATT GAC GCC AGA GC | CARC1 (pericentric; 3 F) |
| TE311 | AAG TAC GCG TAC GAA GCA TC | CARC1 (pericentric; 3 R) |
| TE306 | CAA ACC ACC TCT TAC GTC GTT G | CARC1 (pericentric; 4F) |
| TE307 | TTT CGT GCA CTG CGT TCA AG | CARC1 (pericentric; 4R) |
| TE308 | TCC TGG AAT GGA GAC CGT TTT C | CARC1 (pericentric; 5F) |
| TE309 | AGC CGA CAA ATT TCG TGC AC | CARC1 (pericentric; 5R) |
| TE373/MSB180 | ACT TTG GTT TTC CGG TGT GC | CARC1 (pericentric; 6F) |
| TE374/MSB181 | CCA GCG ATG AGA TGC GAA AAG | CARC1 (pericentric; 6R) |
| TE377 | TCG CTT TTC GCA TCT CAT CG | CARC1 (pericentric; 9F) |
| TE378 | AGC GGG CGG GTT ATA AAT AAC | CARC1 (pericentric; 9R) |
| TE533 | ACC TTC TAC TTC CAT GCC GTT G | CARC1 (pericentric; 10F) |
| TE534 | TGC GTG CCG ATG TAG AAT TG | CARC1 (pericentric; 10R) |
